# Neuronal activity in the human amygdala and hippocampus enhances emotional memory encoding

**DOI:** 10.1101/2021.11.28.470278

**Authors:** Salman E. Qasim, Uma R. Mohan, Joel M. Stein, Joshua Jacobs

## Abstract

Emotional events comprise our strongest and most valuable memories, yet it is unknown how the brain prioritizes emotional information for storage. Here, we examined the neural basis of this prioritization using direct brain recording, deep brain stimulation, and psychometric assessment, with human subjects performing an episodic memory task in which they showed improved performance for emotional stimuli. During the task, high-frequency activity (HFA), a correlate of neuronal spiking activity, increased in both the hippocampus and amygdala when subjects successfully encoded emotionally arousing stimuli. Applying inhibitory electrical stimulation to these regions decreased HFA and specifically reversed the enhancement of memory for emotional stimuli, indicating that neuronal activity in the amygdalohippocampal circuit has a direct role in prioritizing emotional memories. Finally, we found abnormal patterns of amygdalohippocampal HFA in depressed individuals which correlated with a bias for negative memories in these subjects. Going forward, targeted modulation that upregulates neuronal excitation in the amygdalohippocampal circuit may have a causal and translational role in modulating emotional memory.

## Introduction

We remember emotional events better than neutral ones^1–3^. This enhanced recollection for emotional information is important practically for protecting our most important memories, and may also provide generalizable clues about the fundamental nature of memory^4^, by explaining how the brain remembers some events better than others. One of the critical brain regions for processing emotional stimuli^5,6^—the amygdala—is an early target of Alzheimer’s disease^7^ and abuts the anterior portion of the hippocampus, the brain region most strongly associated with declarative memory^8^. This etiological and anatomical proximity converges with behavioral, imaging, and lesion evidence that the amygdala may be critical for memory of emotional events^9–17^.

One prominent theory of the amygdala’s role in memory proposes that the amygdala boosts hippocampal encoding and consolidation of emotional stimuli by facilitating the release of norepinephrine from the locus coeruleus (LC)^18,19^. While it is difficult to directly measure human norepinephrine fluctuations, there is indirect evidence for this theory from pharmacological studies showing that enhancing^20,21^ or disrupting^22,23^ noradrenergic transmission, respectively, enhances and impairs memory for arousing stimuli. Noradrenergic inputs may modulate the amygdalohippocampal circuit by up-regulating the mean rate of neuronal activity, as suggested by both direct recordings of neuronal activity^24–26^ as well as recordings of high-frequency activity (HFA) in limbic local field potentials^27,28^. Similarly, data from patients with depression also show links between emotional memory, norepinephrine, and amygdala activity. Depressed individuals, who exhibit impaired emotional memory^29^, show improvement in depressive symptoms when treated by norepinephrine agonists^30^ or when receiving brain stimulation that increases amygdala HFA^31^. Together, these findings suggest that HFA in the hippocampus and amygdala is, at least in part, driven by noradrenergic up-regulation of neural activity^32^.

Building off these ideas, here we hypothesized that our ability to prioritize emotionally salient information for improved memory would rely on this up-regulation of neuronal activity within the amygdala and hippocampus during encoding. We tested this hypothesis in humans using a novel tripartite approach that combined three complementary methods for the first time: direct human brain recordings, deep brain stimulation, and psychometric assessment of depressive symptoms in epilepsy patients performing a verbal free recall task. Free recall is an episodic memory task in which subjects exhibit enhanced memory for emotional words^33–35^, and elicits norepinephrine release during successful memory encoding^36^. Thus, we hypothesized that direct brain recordings from subjects performing this task would reveal if HFA reflected noradrenergic dynamics during the prioritized encoding of emotional events. Consistent with these predictions, we found that the amplitude of HFA in the amygdala and hippocampus increased with the encoding of emotionally arousing words^37,38^. Inversely, we found that inhibitory brain stimulation weakened HFA and impaired recall of emotional words, suggesting that there is a causal relationship in the amygdalohippocampal circuit between neuronal spiking and the enhanced memory for emotional items. Finally, we demonstrated that depressed subjects—whose impaired emotional processing is characterized by disruption of noradrenergic neurotransmission^39^—exhibited abnormal patterns of amygdalohippocampal HFA during emotional memory encoding that correlated with their bias towards negative memories compared to non-depressed individuals. Overall, our findings demonstrate that neuronal activity in the human amygdala and hippocampus, a potential correlate of noradrenergic up-regulation, causally supports the prioritization of emotional memories.

## Results

### Emotional stimuli are better remembered

We analyzed data from 165 subjects who performed a verbal episodic memory task where they viewed and remembered lists of words. After each list, subjects performed a math distractor task to prevent rehearsal, and were then told to recall as many words as possible, in any order (Fig. 1A). We quantified the valence and arousal of each word using a publicly available lexicon. This allowed us to assess whether the emotional properties of each word impacted subject’s memory encoding (see Methods for detail)^40^. The words in our dataset exhibited higher valence (0.56±0.15 a.u.) and lower arousal (0.36±0.16 a.u.) than the broader lexicon (p’s< 3*x*10^-9^, Wilcoxon rank-sum test, Fig. 1B). Subjects recalled 26% of words on average (Fig. 1D), and arousal and valence ratings modulated recall performance, with subjects better remembering words that were associated with high arousal and/or negative valence (Fig. 1E; S1). Arousal and valence also modulated recall latency (Fig. 1F) and clustering during recall (Fig. S2).

**Figure 1:**
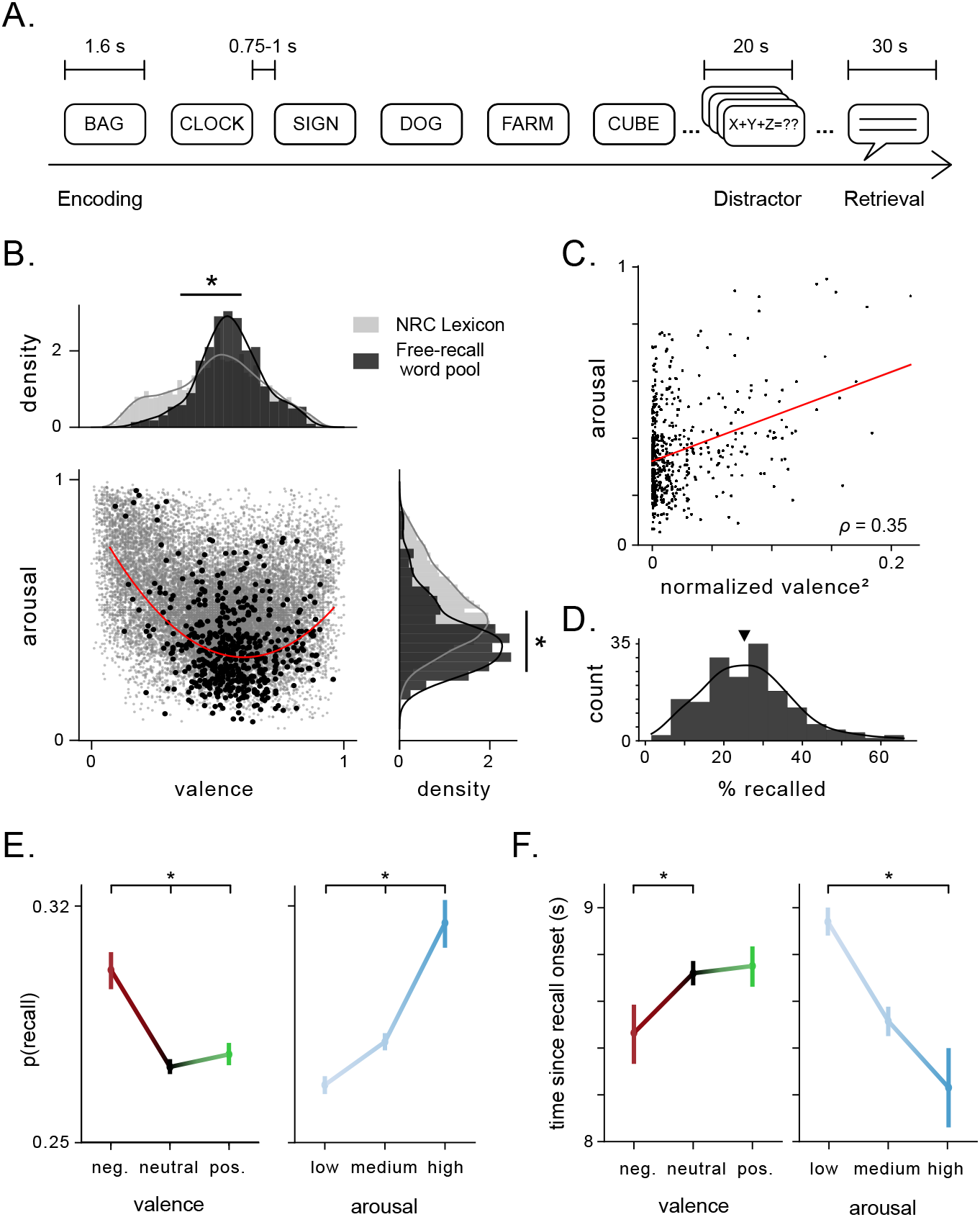
Emotional features of stimuli in a verbal free recall task influence recall performance. A) Schematic of task design showing the time intervals during and between task stages. Subject encoded 12 words per list. B) Joint scatter plot and marginal distributions of valence and arousal ratings in the general rating lexicon (gray) and the word pool for the free recall tasks performed by subjects (black). Asterisks indicate significant difference between the mean ratings for valence and arousal between the word pool and the general lexicon. Red line denotes polynomial fit to valence and arousal ratings of free-recall word pool. C) Arousal plotted as a function of mean-normalized, squared valence. Red line denotes linear fit. Pearson’s correlation coefficient is indicated. D) Distribution of recall performance across all subjects (mean = 26%, black arrow). E) Probability of recall significantly differed as a function of valence (p< 5 × 10^-8^, *χ*^2^(2) = 33.8) and arousal (p< 2 × 10^-15^, *χ*^2^(2) = 68.6). Vertical bars denote standard error. Asterisks denote significant difference in proportions across categories. F) Time since recall onset tended to be shorter for negative vs. neutral words (p= 0.05, *t* = −1.94), and was significantly lower for high vs. low arousal words (p< 4 × 10^-5^, *t* = −4.16). Asterisks denote significant difference in recall times across categories.

The emotional features of each word were thus predictive of subjects’ memory performance, even as the task did not explicitly depend on the emotional features of the words, consistent with earlier work^33,34^. These behavioral results suggest that even when emotional context has limited task relevance, it drives memory enhancement and is thus likely predictive of subjects’ implicit emotional responses. Because the amygdala is broadly involved in the enhancement of memory for emotional events^9,12–14^, and delayed free recall tasks depend on the hippocampus^41^, we hypothesized that the joint activation of amygdala and hippocampus was responsible for this phenomenon.

### Neuronal activity in the hippocampus and amygdala scales with word arousal during successful encoding

We next tested how neuronal activity in the hippocampus and amygdala corresponded to the effects of emotional context on memory. We analyzed brain recordings from intracranial electroencephalographic (iEEG) electrodes (n=965 in the hippocampus, n=411 electrodes in the amygdala) implanted in these subjects while undergoing intracranial monitoring for epilepsy treatment (Fig. 2A). A majority of amygdala electrodes were located in the basolateral nuclei (Fig. S3). First, we assessed whether spectral power of the signals at each electrode during encoding was predictive of subsequent recall by comparing the power spectrum of the iEEG signals during encoding between remembered vs. forgotten items^42,43^. Prior iEEG studies have identified a characteristic increase in high-frequency power and decrease in low-frequency power in the hippocampus associated with encoding of successfully recalled words^44–47^, which we replicated. In addition, we also demonstrated that the amygdala showed similar iEEG power increases during encoding of subsequently remembered items in the HFA band (30–128 Hz) and a more muted decrease in theta/alpha (2–12 Hz) power (Figs. 2B, S4)). These effects were not a result of differences in spectral tilt or changes to the height or frequency of the spectral peaks (Fig. S5). These patterns appeared to be lateralized to the left hemisphere (Fig. S6). Overall, these results thus suggests that increased HFA power in the amygdala and hippocampus predict successful memory encoding.

**Figure 2:**
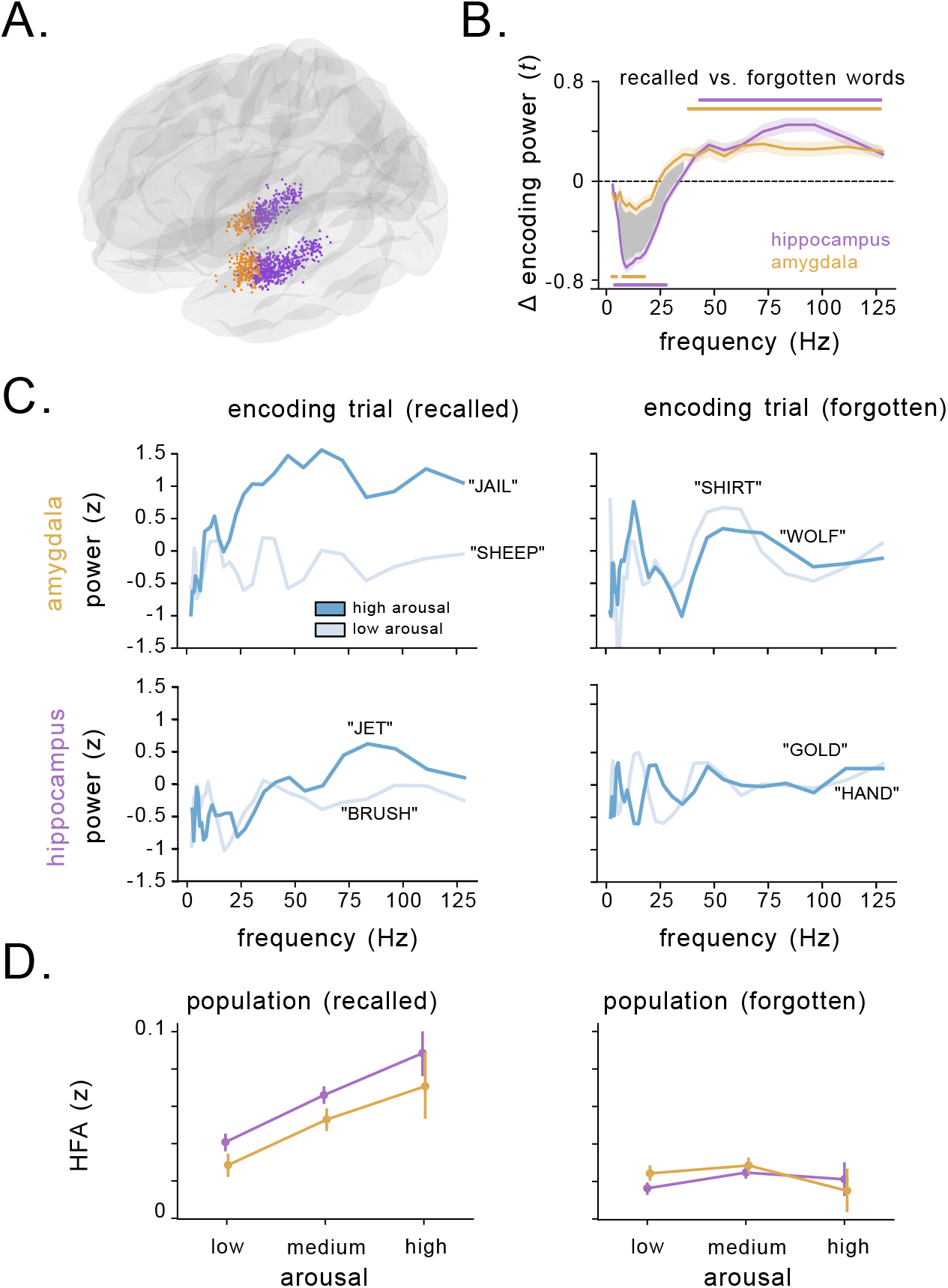
Arousal modulates high-frequency activity related to successful memory encoding in the hippocampus and amygdala. A) Location of all 1,376 electrodes recorded across all subjects. Purple circles indicate electrodes localized to hippocampus, orange circles indicate electrodes localized to the amygdala. B) Comparison of SME (t-statistic) averaged over hippocampal (purple) and amygdala (orange) electrodes. Horizontal lines denote SMEs that significantly deviate from 0 for electrodes in the hippocampus (p’s<= 0.001, cluster-permutation test), and the amygdala (p’s<= 0.02, cluster-permutation test). The shaded region denotes a significant difference between the magnitude of the SME for hippocampus vs. amygdala (p’s<= 0.001, cluster-permutation test). C) Within-list z-scored power during example encoding trials from a single amygdala (top) and hippocampal (bottom) electrode in two subjects during memory encoding. Power during encoding of a high arousal word from the list is depicted in dark blue, while power for a low arousal word from the list is depicted in light blue. HFA increases during successful encoding (left) of high arousal words vs. low arousal words, but not during failed encoding (right). D) Z-scored HFA in the amygdala (orange) and hippocampus (purple) during the encoding phase as a function of word arousal for recalled (left) and forgotten (right) words. Circles represent mean of binned arousal, with vertical lines denoting the standard error of the bin.

We next tested whether these memory-related spectral dynamics were mediated by the emotional features of each word, as we hypothesized that the signals might reflect up-regulation of neuronal spiking caused by noradrenergic release related to emotional processing. Figure 2C shows an example of the signals we observed in the hippocampus and amygdala, depicting z-scored power spectra from individual trials when subjects viewed words. For words that were subsequently recalled (left panel), HFA was higher for more arousing words. This arousal-related HFA effect was not present for words that subjects would subsequently forget (right). To assess these effects statistically, we used a mixed-effects linear model to assess how spectral power during encoding changed as a function of valence, arousal, memory performance, brain region, and hemisphere (see Methods). Consistent with the examples shown above, HFA was significantly modulated by the interaction of arousal and memory performance (*χ*_(15)_ = 31.3, p= 0.008, likelihood ratio test). This pattern is illustrated in Figure 2D: over the entire dataset, HFA in the hippocampus and amygdala exhibited a clear increase with arousal for remembered (left), but not forgotten words (right). These effects were consistent across both hemispheres. In contrast, while theta/alpha power changes varied with brain region and memory performance (*χ*_(16)_ = 29.9, p= 0.018, likelihood ratio test), they were not significantly modulated by word arousal or valence (*χ*_(15)_ = 12.4,16.4, p’s> 0.3, likelihood ratio test). The link between HFA and arousal for successfully encoded words was consistent even at the level of individual words grouped across subjects, with high arousal words showing larger HFA contrasts between remembered and forgotten trials compared to low arousal words (Fig. S7A, B). Critically, though arousal and valence were correlated (Fig. 1C), valence was not a significant modulating factor in the model (Fig. S8), consistent with the notion that noradrenergic modulation of memory is independent of valence^48^. The increase in HFA was not accompanied by significant changes in high-frequency coherence or cross-frequency coupling between the amygdala and hippocampus (p’s> 0.05, cluster-permutation test). These results thus demonstrate that word arousal specifically up-regulated amygdalohippocampal HFA during encoding of subsequently remembered words.

### Electrical stimulation of the amygdalohippocampal circuit selectively disrupts HFA and memory for emotional stimuli

To better understand whether the HFA we observed reflects causal neural mechanisms, we next tested whether disrupting neuronal activity in the amygdalohippocampal circuit would impair memory for emotional information. To do so, we analyzed the effect of high-frequency (50 Hz) deep brain stimulation on memory performance. This type of stimulation has been shown to modulate HFA^49^ and impair memory when applied to medial temporal brain regions^50^. A group of 19 subjects performed 32 sessions of the free recall task while direct electrical stimulation was applied to their hippocampus and amygdala (Table 2, Fig. S9, 3A, see *Methods*). In addition, 8 subjects were stimulated in non-hippocampal MTL regions such as parahippocampal gyrus and perirhinal cortex, which served as nearby control regions outside of the amygdalohippocampal circuit.

**Figure 3:**
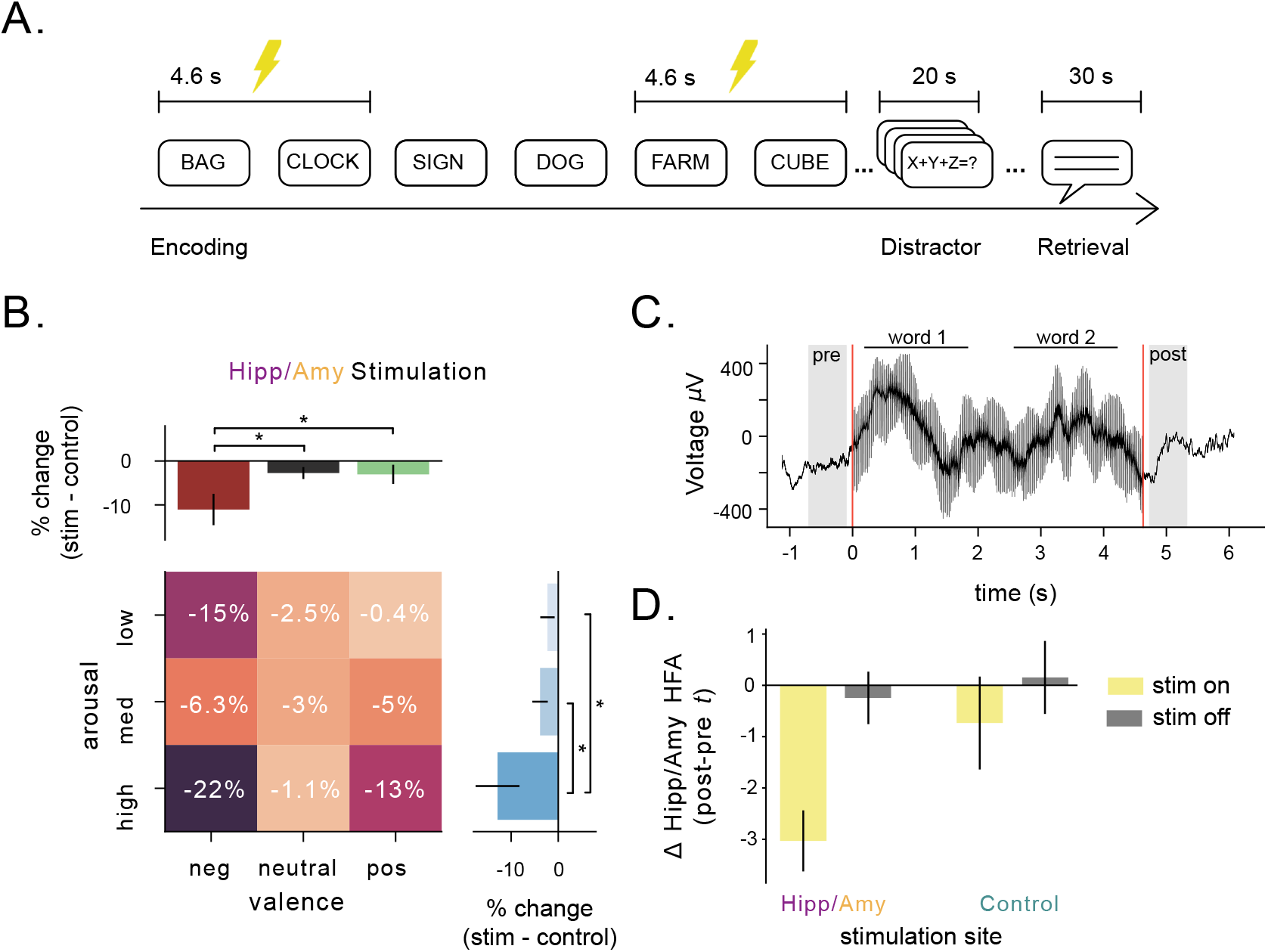
Direct stimulation of hippocampus and amygdala during encoding impairs emotion-mediated memory and decreases HFA. A) Schematic of task design showing the time intervals during direct brain stimulation. Stimulation was applied to alternating pairs of words in a list. B) Effect of stimulation administered to electrodes located either in the hippocampus or the amygdala, split by arousal and valence. Asterisks indicate significant differences in the effect of stimulation on recall as a function of emotional features of words (p’s< 0.05, likelihood-ratio test). Heatmap numbers indicate percentage of change in recall performance during stimulation. C) Single-trial example local-field potential during stimulation of a word-pair. Shaded regions indicate pre and post-periods used for analysis. Word presentation is indicated by horizontal lines. Red lines indicate onset and offset of stimulation. D) The difference in HFA in the hippocampus and amygdala (z-scored) before and after stimulation, measured by a paired t-test. The yellow bars denote words with stimulation turned on, while gray bars denote words with stimulation turned off. Bars on the left side correspond to stimulation applied to the hippocampus or amygdala, while bars on the right side correspond to stimulation applied to control regions in the MTL.

**Table 1:** Subject demographics. Age, gender, and BDI-II score (where available) for the subjects that participated in the verbal free recall task.

| subject | gender | age | BDI-II | subject | gender | age | BDI-II | subject | gender | age | BDI-II |
| --- | --- | --- | --- | --- | --- | --- | --- | --- | --- | --- | --- |
| R1001P | Female | 48 | 2 | R1122E | Female | 48 | - | R1240T | Female | 38 | - |
| R1002P | Female | 49 | 0 | R1123C | Female | 29 | 19 | R1241J | Male | 52 | 16 |
| R1003P | Female | 39 | - | R1124J | Female | 41 | 4 | R1243T | Male | 64 | - |
| R1004D | Female | 52 | - | R1125T | Female | 45 | - | R1245E | Male | 30 | - |
| R1006P | Female | 21 | 7 | R1128E | Male | 26 | - | R1247P | Female | 62 | 0 |
| R1010J | Female | 31 | 1 | R1131M | Male | 24 | - | R1254E | Male | 29 | 17 |
| R1015J | Female | 55 | 15 | R1134T | Male | 65 | - | R1260D | Female | 57 | 12 |
| R1020J | Female | 49 | 4 | R1136N | Female | 57 | - | R1261P | Female | 47 | - |
| R1022J | Male | 25 | 1 | R1137E | Female | 21 | - | R1266J | Female | 48 | 41 |
| R1023J | Male | 33 | 2 | R1138T | Male | 42 | - | R1268T | Female | 33 | - |
| R1027J | Male | 48 | 19 | R1144E | Male | 53 | - | R1269E | Female | 42 | - |
| R1028M | Female | 27 | - | R1145J | Male | 46 | 5 | R1271P | Male | 37 | - |
| R1030J | Male | 23 | 11 | R1147P | Male | 48 | 4 | R1273D | Female | 48 | 18 |
| R1031M | Male | 40 | - | R1148P | Female | 60 | 0 | R1275D | Male | 42 | 8 |
| R1032D | Female | 20 | 6 | R1150J | Female | 50 | 18 | R1278E | Female | 22 | - |
| R1033D | Female | 32 | 3 | R1151E | Male | 36 | - | R1279P | Female | 58 | 7 |
| R1034D | Female | 29 | 10 | R1153T | Male | 38 | - | R1281E | Female | 44 | - |
| R1035M | Female | 45 | 10 | R1154D | Female | 36 | 31 | R1283T | Female | 29 | - |
| R1044J | Male | 59 | 7 | R1157C | Male | 22 | - | R1286J | Female | 58 | 16 |
| R1045E | Male | 51 | - | R1158T | Female | 45 | - | R1288P | Male | 53 | - |
| R1048E | Male | 22 | 7 | R1159P | Female | 43 | 7 | R1292E | Female | 39 | 27 |
| R1049J | Female | 52 | 6 | R1161E | Female | 54 | - | R1293P | Male | 38 | - |
| R1052E | Female | 20 | - | R1162N | Female | 31 | - | R1297T | Male | 25 | - |
| R1053M | Female | 39 | - | R1163T | Male | 46 | - | R1298E | Female | 25 | - |
| R1054J | Male | 24 | 1 | R1164E | Male | 38 | - | R1299T | Male | 44 | - |
| R1056M | Male | 34 | - | R1167M | Male | 33 | - | R1303E | Female | 63 | - |
| R1059J | Female | 45 | 13 | R1168T | Male | 24 | - | R1306E | Male | 46 | - |
| R1061T | Male | 21 | - | R1169P | Female | 43 | 37 | R1308T | Female | 40 | - |
| R1062J | Female | 24 | 9 | R1171M | Male | 36 | - | R1309M | Female | 22 | - |
| R1063C | Male | 24 | - | R1172E | Female | 22 | - | R1310J | Male | 20 | 0 |
| R1065J | Female | 34 | 8 | R1174T | Male | 29 | - | R1311T | Male | 22 | - |
| R1066P | Male | 39 | 1 | R1176M | Female | 41 | - | R1313J | Male | 22 | 1 |
| R1067P | Female | 45 | 1 | R1180C | Female | 21 | - | R1315T | Male | 22 | - |
| R1068J | Female | 39 | 7 | R1186P | Male | 28 | - | R1316T | Female | 33 | - |
| R1077T | Female | 48 | 21 | R1188C | Female | 26 | - | R1320D | Male | 57 | - |
| R1080E | Female | 43 | 5 | R1190P | Female | 58 | 8 | R1323T | Male | 40 | - |
| R1083J | Female | 49 | 19 | R1191J | Male | 19 | - | R1325C | Female | 51 | - |
| R1086M | Male | 21 | - | R1192C | Male | 28 | - | R1328E | Female | 28 | - |
| R1089P | Male | 36 | - | R1195E | Male | 44 | - | R1330D | Female | 38 | 8 |
| R1092J | Male | 45 | 13 | R1200T | Male | 26 | - | R1331T | Female | 52 | - |
| R1094T | Male | 47 | - | R1201P | Male | 37 | 1 | R1332M | Male | 27 | - |
| R1096E | Female | 34 | - | R1203T | Female | 36 | - | R1334T | Female | 38 | - |
| R1100D | Female | 44 | 39 | R1204T | Female | 26 | - | R1336T | Female | 51 | - |
| R1101T | Female | 26 | - | R1212P | Male | 47 | - | R1337E | Male | 55 | - |
| R1102P | Male | 35 | - | R1215M | Female | 51 | - | R1338T | Male | 22 | - |
| R1104D | Male | 22 | - | R1216E | Female | 23 | - | R1339D | Male | 29 | 5 |
| R1105E | Male | 25 | 0 | R1217T | Male | 37 | - | R1341T | Male | 33 | - |
| R1106M | Male | 27 | - | R1221P | Male | 57 | 0 | R1342M | Male | 42 | - |
| R1108J | Female | 24 | 1 | R1226D | Female | 42 | 16 | R1343J | Male | 35 | 1 |
| R1112M | Female | 30 | - | R1227T | Male | 33 | - | R1346T | Female | 38 | - |
| R1113T | Female | 37 | - | R1228M | Female | 59 | - | R1349T | Female | 44 | - |
| R1114C | Female | 32 | - | R1229M | Male | 29 | - | R1350D | Male | 50 | 20 |
| R1115T | Male | 47 | - | R1230J | Female | 56 | 24 | R1374T | Male | 60 | - |
| R1119P | Female | 27 | 9 | R1236J | Female | 52 | 14 | R1397D | Female | 34 | 11 |
| R1120E | Female | 34 | - | R1239E | Male | 27 | 19 | R1409D | Female | 49 | - |

**Table 2:**
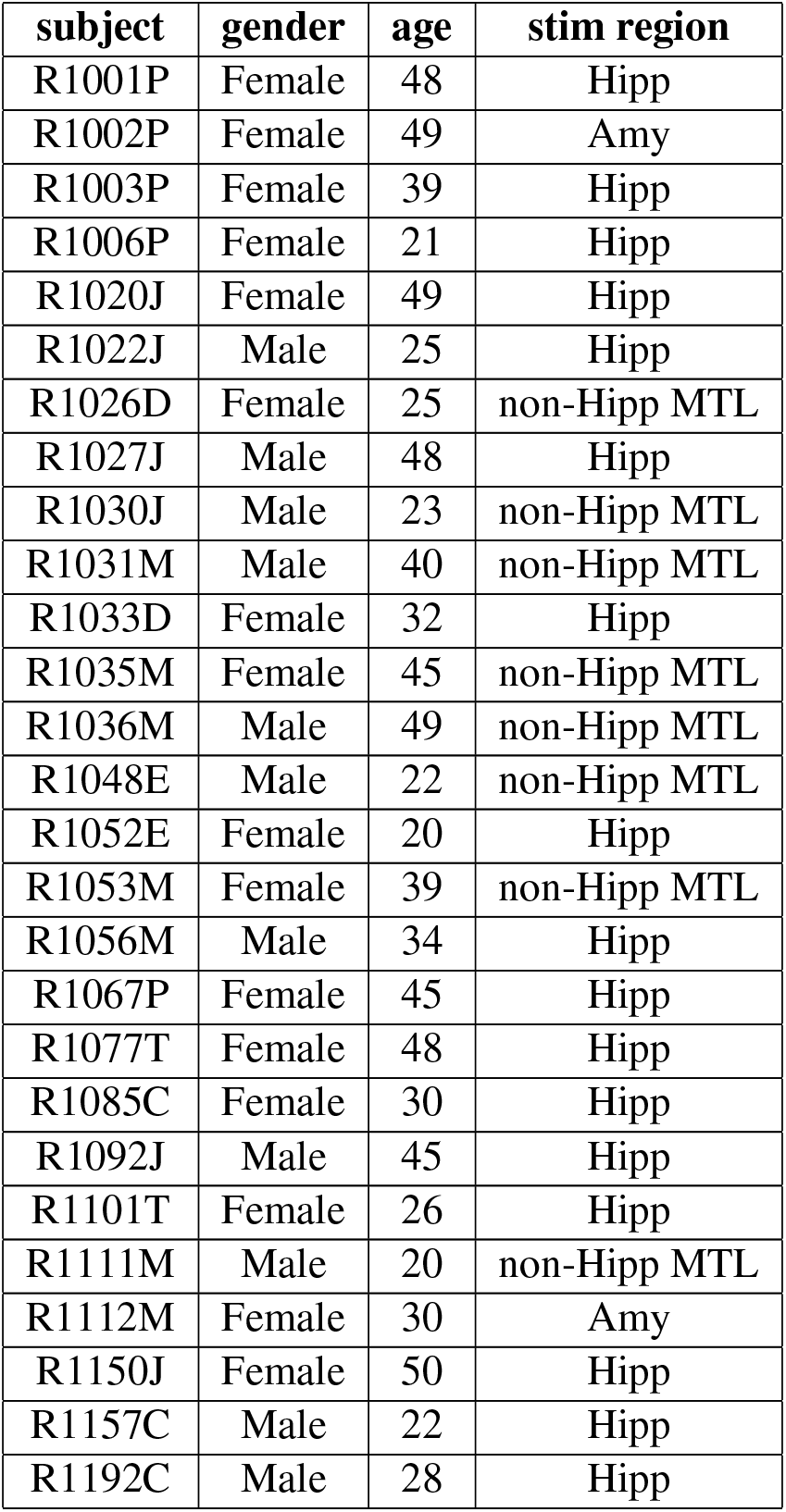
Subject demographics for stimulation task. Age, gender, and region of stimulating electrodes for the subjects that participated in the stimulation version of the verbal free recall task.

| **subject** | **gender** | **age** | **stim region** |
| --- | --- | --- | --- |
| R1001P | Female | 48 | Hipp |
| R1002P | Female | 49 | Amy |
| R1003P | Female | 39 | Hipp |
| R1006P | Female | 21 | Hipp |
| R1020J | Female | 49 | Hipp |
| R1022J | Male | 25 | Hipp |
| R1026D | Female | 25 | non-Hipp MTL |
| R1027J | Male | 48 | Hipp |
| R1030J | Male | 23 | non-Hipp MTL |
| R1031M | Male | 40 | non-Hipp MTL |
| R1033D | Female | 32 | Hipp |
| R1035M | Female | 45 | non-Hipp MTL |
| R1036M | Male | 49 | non-Hipp MTL |
| R1048E | Male | 22 | non-Hipp MTL |
| R1052E | Female | 20 | Hipp |
| R1053M | Female | 39 | non-Hipp MTL |
| R1056M | Male | 34 | Hipp |
| R1067P | Female | 45 | Hipp |
| R1077T | Female | 48 | Hipp |
| R1085C | Female | 30 | Hipp |
| R1092J | Male | 45 | Hipp |
| R1101T | Female | 26 | Hipp |
| R1111M | Male | 20 | non-Hipp MTL |
| R1112M | Female | 30 | Amy |
| R1150J | Female | 50 | Hipp |
| R1157C | Male | 22 | Hipp |
| R1192C | Male | 28 | Hipp |

We analyzed these data to test how stimulation in the amygdala and hippocampus impacted memory for words of varying arousal and valence. Figure 3B shows that high-frequency stimulation, when applied to the hippocampus or amygdala, specifically impaired encoding for emotional words. Specifically, recall of both high arousal words and negative words was significantly more impaired by stimulation than other categories of words (p’s< 0.05, likelihood-ratio tests). This was not the case for stimulation in control regions (Fig. S10A), suggesting that hippocampal/amygdalar stimulation was selectively disrupting neural dynamics related to the emotional enhancement of memory. To explain how stimulation specifically modulated the encoding of emotionally relevant stimuli, we compared spectral power in the amygdalohippocampal circuit pre versus post stimulation (Fig. 3C). Stimulating this circuit caused a significant reduction in amygdalohippocampal HFA (*t* = −4.84, p< 0.001, one-sample t-test), and this effect was significantly weaker when stimulation was applied to adjacent control regions (F= −5.6, p= 0.02, ANOVA). Stimulation did not significantly decrease theta power in the hippocampus and amygdala (F= 2.57, p= 0.12, ANOVA), suggesting that amygdalohippocampal HFA, specifically, underlies the stimulation-induced disruption of emotional memory enhancement.

### Depression disrupts arousal-mediated memory and amygdala–hippocampus HFA

A prominent theory of affective disorders hypothesizes that deficiencies in noradrenergic neurotransmission underlie depressive symptoms, including the tendency to ruminate more on negative than positive memories^39^. Therefore, we next assessed whether depression-elicited changes in amygdalar and hippocampal HFA corresponded to a bias towards negative memories. We examined this by comparing HFA signals between subjects who did and did not show evidence of depression. We had information about depression severity from a subset of the patients in our dataset who had completed the Beck Depression Inventory (BDI-II). There was a range of depression levels across our dataset (Fig. 4A). As expected from earlier work^51^, depressed subjects exhibited worse memory performance (Fig. 4B), *t* = −2.4, p=0.02). We next examined how the memory encoding from depressed individuals correlated with the emotional features of each to-be-remembered word. Notably, depressed subjects exhibited a weaker link between arousal (how engaging a word is) and memory performance compared to subjects without depression (*χ*^2^ = 4.7, 15.3, respectively, Fig. 4C). Conversely, depressed subjects demonstrated a larger dependence on valence (whether a word is positive or negative) than non-depressed subjects (*χ*^2^ = 23.4,3.9, respectively, Fig. 4C), with a specific bias for negative memories. These patterns suggest that depressed subjects effectively change strategies during encoding, relying relatively more on word valence during memory encoding, rather than word arousal, in line with the known bias for negative memories in depression^29^.

**Figure 4:**
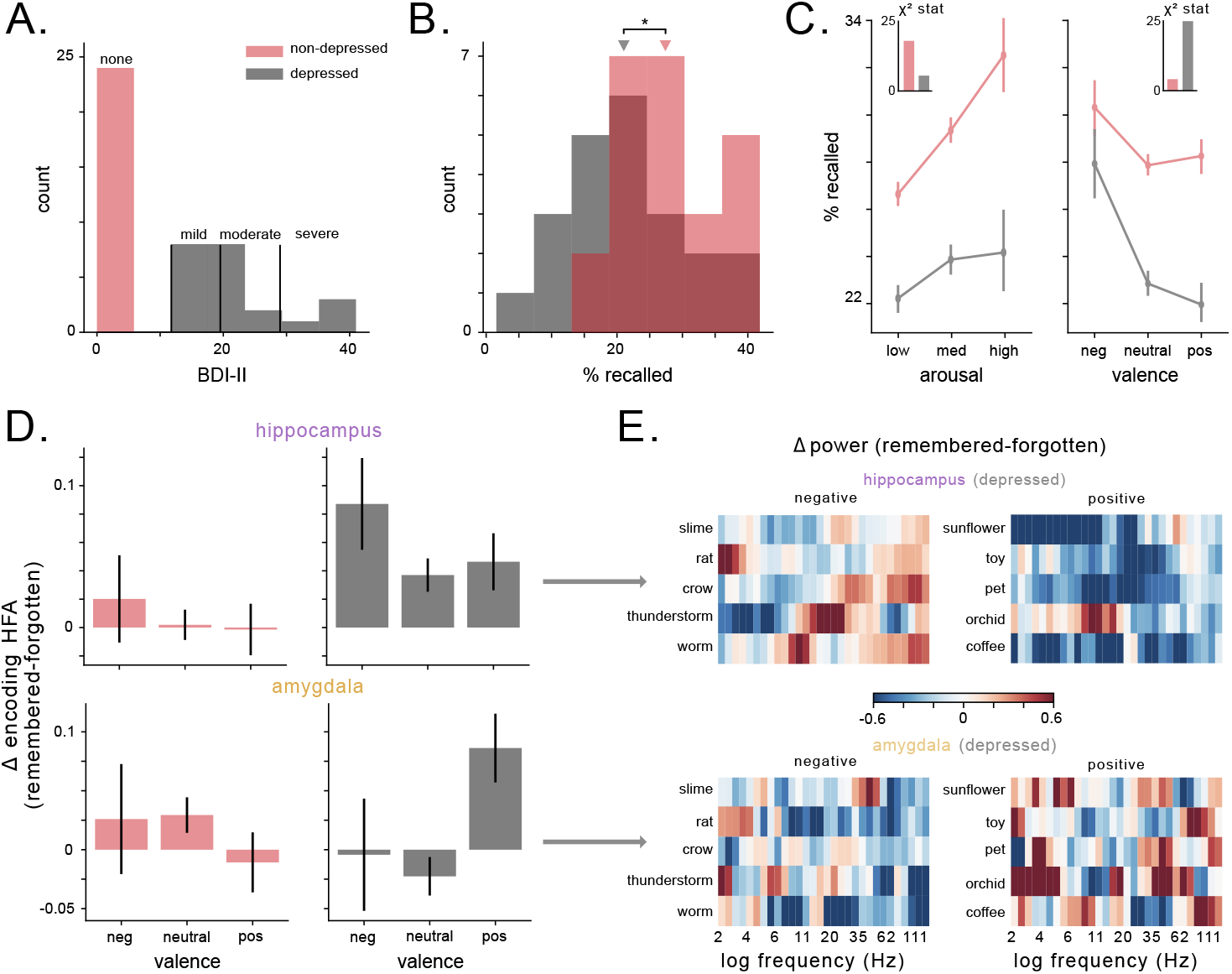
Valence, rather than arousal, modulates memory performance and HFA in subjects with depression. A) Histogram of BDI-II scores for patients, split by depressive characterization. B) Distribution of recall performance across subjects as a function of depression rating. Depressed patients exhibit significantly worse recall performance (*t* = −2.4, *p* = 0.02). C) Recall percentage expressed as a function of arousal (left) and valence (right) for both depressed and non-depressed subjects. Inset depicts the *χ*-statistic assessing the affect of arousal and valence on recall performance in both groups of subjects. D) The difference in HFA (z-scored) during the encoding period for remembered and forgotten items, for subjects with and without depression in the hippocampus (top) and amygdala (bottom). E) Heatmaps of hippocampal (top) and amygdalar (bottom) power (z-scored within session) for specific words from the task wordpool, averaged across sessions and subjects. Words were selected from the 30 words with the lowest valence ratings (left) or highest valence ratings (right). Warm colors indicate higher values while cool colors indicate lower values.

Subjects with depression also show abnormal memory-related neural activity in the amygdalohippocampal circuit. First, depressed subjects showed a broadband enhancement of memory-related spectral power changes compared to non-depressed individuals (*χ*_(4)_ = 10.1, 14, p’s< 0.04, Fig. S11). Critically, mixed-effects models revealed that valence significantly predicted changes to memory-related HFA, while arousal did not (*χ*_(15)_ = 17.9, p= 0.21), in concordance with behavioral shift from arousal-to valence-mediated memory in depressed subjects. This valence-driven pattern of HFA during successful memory encoding varied with depression severity and brain region (*χ*_(15)_ = 40.5, p< 4*x*10^-4^, likelihood ratio test). Specifically, when compared to non-depressed subjects, depressed subjects exhibited elevated hippocampal (but not amygdalar) HFA while encoding negative words, and elevated amygdalar (but not hippocampal) HFA while encoding positive words (Fig. 4D). Figure 4E depicts examples of this effect in depressed subjects at the level of individual positive and negative words. By demonstrating that depression elicits distinct valence-driven HFA patterns in the amygdala and hippocampus in congruence with a valence-driven shift in memory recall performance, these results suggest that HFA in the amygdalohippocampal circuit may underlie the valence-driven rumination that occurs in depression.

## Discussion

The emotional context of an event often determines how that event is remembered. Here, we investigate the neural basis of our enhanced memory for emotional events by using direct recordings and stimulation in human neurosurgical subjects. By directly examining human hippocampal and amygdalar electrophysiology from subjects performing a memory task, we assessed the role of these two brain structures in encoding memories with emotional associations. In both the hippocampus and amygdala we found that high-frequency activity (HFA), a proxy for local neuronal spiking, correlated with stimulus-induced arousal during successful memory encoding. We found that this phenomenon is causally important because perturbing this network with disruptive brain stimulation selectively impaired the recall of emotional stimuli. Finally, we found that compared to normal individuals, depressed subjects prioritize valence information over arousal during recall, in concordance with altered, valence-driven amygdalohippocampal HFA. These results (1) demonstrate that up-regulation of amygdalohippocampal activity during encoding is correlated with enhancement of memory for emotionally engaging stimuli in humans, and (2) show that modulating the activity within this circuit causally affects how the human brain prioritizes certain information for memory encoding.

Substantial behavioral evidence has shown that the brain prioritizes the encoding of emotional content^1,2,33,34^. While lesion studies have demonstrated the importance of both the amygdala and hippocampus to this behavioral effect, our findings show a causal mechanism underlying this effect: increased neuronal activity in the amygdalohippocampal circuit, as indexed by HFA, enhances memory for emotional information during memory encoding. This finding bridges evidence from iEEG studies of memory that implicated increased HFA with successful word recall^44, 46^ and aversive image viewing^52^ with fMRI studies that have demonstrated increasing activation in the amygdala and hippocampus with recall of more emotional stimuli^12,15,53^. Furthermore, the results from our stimulation experiments indicate that this amygdalohippocampal up-regulation is causally responsible for the emotional enhancement of memory, because decreasing HFA in these structures with stimulation reversed the memory enhancement that was caused by emotional context.

While our study is the first, to our knowledge, that examines the effects of direct stimulation of the amygdala–hippocampus system on affective memory processes specifically, two recent studies are related to ours because they separately probed the effect of amygdala stimulation on memory and affective disorders. Inman et al. showed that amygdala stimulation enhanced memory overall, in contrast to our observed pattern of stimulation-induced memory impairment and HFA decreases^54^. Three key methodological differences between that study and our work may explain the differing results. First, the stimuli used by Inman et al., were neutral stimuli, which likely did not engage the same emotional memory processes as the stimuli in our task. Second, whereas in our study retrieval occurred a few minutes after encoding, Inman et al. observed memory enhancement one day after stimulation, suggesting that amygdala stimulation may have affected later memory consolidation processes rather than strictly modifying memory encoding. The second study assessed how amygdala stimulation modulated human depression symptoms, showing that amygdala HFA was a successful biomarker for high-efficacy closed-loop stimulation to treat major depressive disorder in a single subject^31^. Not only was bilateral amygdala HFA sufficient to classify depressive states, but stimulation induced a reduction in HFA that improved symptom severity. Our work provides a potential mechanistic explanation for these stimulation results by demonstrating that amygdalar and hippocampal HFA correlate with memory for emotional stimuli. Future work should thus establish whether amygdala stimulation that reduces depressive symptoms also modulates memory for emotional content.

Examining the link between emotional state and memory is particularly important given our results showing a direct relation between HFA, depression, and the emotional enhancement of memory. We found that depressed subjects, in contrast to non-depressed individuals, remembered words better on the basis of their valence rather than their arousal. This is potentially important because one prominent theory^55^ suggested that valence-mediated memory is thought to engage prefrontal–hippocampal pathways, in contrast to arousal-mediated memory, which is thought to rely more heavily on amygdalohippocampal interaction. Thus this hypothesis, with our findings, supports the idea that amygdalar involvement in memory may be disrupted in subjects with depression. Consistent with this idea, the enhanced memory that depressed subjects exhibited for negative words only increases HFA in the hippocampus, not the amygdala. Meanwhile, the amygdala increase in HFA corresponded with the recall of words with positive affect, which is the specific word category at which depressed subjects performed worst. There are several findings from the literature that are consistent with this interpretation. Electrical stimulation of the amygdala induces increased bias for positive affect^56^. Furthermore, depressed subjects’ impairment in positive stimulus recall can be rescued through administration of reboxetine, a norepinephrine agonist^57^. Our data thus suggest that disrupting amygdala-mediated noradrenergic transmission is responsible for the the differences we observed in both memory performance and HFA in depressed subjects, in line with theories of depressive pathology^39, 58^. Overall, our findings implicate HFA within the amygdala–hippocampus circuit in the emotional enhancement of memory in healthy individuals as well as its alteration in individuals with affective disorders such as depression.

Although our study did not directly measure neuromodulatory signals, we believe that the emotion-related HFA signals we observed reflects neuromodulatory dynamics in the hippocampus and amygdala—in particular, noradrenergic drive from the locus coeruleus (LC)^32^. Norepinephrine release causes increases in HFA in many brain regions^28,59,60^, including the hippocampus^61^ and amygdala^62^, and has also been linked to sharp-wave ripples (SWRs) in the hippocampus^63^. Consistent with our hypothesis of a link between HFA and norepinephrine, prior studies of human verbal memory have demonstrated that successful memory encoding positively correlated with HFA power in the medial temporal lobe^44, 46, 64^, as well as norepinephrine-related patterns such pupil dilation^36^ and autonomic measures such as heart-rate and skin-conductance^65^. In addition to explaining our HFA findings, noradrenergic transmission might also explain our stimulation results. Work in rodents demonstrated that memory deficits induced by amygdala stimulation are mitigated in the absence of noradrenergic release, suggesting that norepinephrine is critical for amygdala stimulation to modulate memory^66^. How would noradenergic release improve memory? One possibility is that the norepinephrine release facilitates hippocampal spike-timing-dependent plasticity, leading to enhanced memory^32,67–69^. This idea is supported by the fact that norepinephrine release alone is not sufficient to enhance memory unless it also elicits neuronal spiking in the amygdala^70^. Overall, these data support the view that noradrenergic up-regulation of neuronal spiking in the amygdalohippocampal circuit, reflected by increases in HFA, is a generalizable mechanism for the prioritization of information processing in the brain^71^.

Emotional memories are some of the most valuable memories we have, and untangling the neural mechanisms underlying the relative robustness of such memories may prove critical to treating memory disorders^4^. Our work provides a bridge to basic science research in animals, providing new avenues for researchers to link midbrain noradrenergic transmission to electrophysiological correlates of memory in the amygdala and hippocampus, such as the HFA observed here. Furthermore, by demonstrating how activity in the amygdala–hippocampus circuit supports the intersection of human memory and emotion, our findings provide mechanistic support to future therapeutic studies modulating this circuit in order to treat memory^54^ and mood disorders^31^. Overall, our findings suggest that up-regulation of neuronal activity within the amygdalohippocampal circuit during encoding may be a generalizable mechanism for the prioritization of information for memory encoding in humans.

## Methods

### Data recording and participants

Data was recorded from patients undergoing invasive iEEG monitoring in the course of their treatment for drug-resistant epilepsy. Patients were recruited to participate in a multi-center project, with data collected at Thomas Jefferson University Hospital, Mayo Clinic, Hospital of the University of Pennsylvania, Emory University Hospital, University of Texas Southwestern Medical Center, Dartmouth-Hitchcock Medical Center, Columbia University Medical Center, National Institutes of Health, and University of Washington Medical Center. Experimental protocol was approved by the IRB at each institution and informed consent was obtained from each participant. Electrodes were implanted using localized, penetrating depth electrodes (Ad-tech Medical Instruments, WI). Electrodes were spaced 10 mm apart, and data was recorded using either the Nihon Kohden EEG-1200, Natus XLTek EMU 128 or Grass Aura-LTM64. iEEG signals were sampled at either 500, 1,000, or 1,600 Hz and referenced to an intracranial electrode, or a contact on the scalp or mastoid process.

### Statistical analysis and software

We used the scipy, statsmodels and mne packages in Python for statistical analyses. Many statistical analyses were performed at the group level. For these we utilized mixed-effects linear models with subject identity as a random effect. We used likelihood ratio tests to assess the significance of the model after removing interactions and main effects, in order to determine which factors explained a significant amount of the variance in the dependent variable.

### Task

Subjects participated in a delayed free-recall verbal memory task. During this task, a 10 s countdown preceded each list of 12 words, which were presented for 1600 ms each with interstimulus intervals randomly sampled from between 750-1000 ms. Each list was followed by a math distractor task to prevent rehearsal, lasting at least 20 seconds, during which simple math problems were presented until a response was entered or recall began. A visual cue paired with a 800 Hz tone signaled the start of each recall period, and subjects had 30 seconds to verbally recall as many words from the list of 12 words they had just seen, in any order. These vocal responses were recorded and annotated offline to assess recall accuracy. Subjects encoded and recalled 25 lists in each session, and did not see the same list twice across sessions.

Subjects performed one or both versions of this task that differed in the semantic structure of the word lists. The uncategorized version of the task utilizes a word pool of 300 words, constructed by selecting words from the Toronto word pool with intermediate recall performance (after accounting for recall dynamics and clustering effects inherent to free recall)^72^. This word pool was split into lists of 12 words such that the mean pairwise semantic similarity within list was relatively constant across lists. For the categorized free recall task, the word pool was drawn from user-rated semantic categories (using Amazon Mechanical Turk). Words were sequentially presented as categorical pairs (drawn from the same category), and each list consisted of four words drawn from each of three categories. Two pairs drawn from the the same semantic category were never presented consecutively^73^.

### Characterizing emotional context during encoding

We utilized a publicly available rating-scale, the National Research Council (NRC) Lexicon^40^, to quantify the emotional context of the words present in the word pool for each task. We selected this rating scale, as opposed to other commonly used rating scales, because of the higher number of independent raters involved, and the use of split-half reliability testing for all ratings. In sum, 97% of the words tested had ratings in the NRC Lexicon and were analyzed in this study. To ensure that our results were not driven by the choice of rating scale, we replicated our behavioral finding against an alternative rating scale^74^.

### Electrode localization

To localize the depth electrodes to the different subregions of the amygdala, we used the CIT168 atlas^75,76^. Due to the bipolar reference scheme we utilized, we localized the resulting “virtual” electrodes using the averaged MNI coordinates of the two referenced electrodes. Virtual electrodes were labelled according to the nearest subregion in the CIT168 atlas, within a 5 mm diameter (which corresponds to the inter-electrode distance).

### Spectral analysis

All data was band-stop filtered around 60 Hz to minimize line noise, and data were bipolar referenced to eliminate reference channel artifacts and noise^46^. Local field potential data were downsampled to 256 Hz for analyses. Prior to analysis we removed trials with interictal epileptiform discharges (IEDs) using established criteria depending on frequency, amplitude, and duration parameters^77^. We used a continuous wavelet transform (Morlet wavelets, wave number=6) with 30 log-spaced frequencies between 2 and 128 Hz, and 1000 ms buffer windows to attenuate convolution edge effects. We then averaged power during into two bands: theta (2-12 Hz), and HFA (30-128 Hz).

To assess the putative effect of spectral tilt, peak height, and peak frequency, we utilized the FOOOF algorithm for parameterizing power spectra^78^. For the encoding phase, we computed spectral power during 1500 ms epochs during which the word was visible, and fit a mixed-effects linear model with emotional context, brain region, hemisphere, and subsequent memory as fixed effects, subject identity as a random effect, and oscillatory power in each band (z-scored within-subject) as the dependent variable.

### Spectral connectivity

In order to compute the normalized coherence between the amygdala and hippocampus we generated surrogate timeseries by swapping time blocks and then z-scored the true coherence estimate relative to the coherence computed for this surrogate distribution. We computed the phase-amplitude coupling (PAC) using the modulation index method^79^. We also normalized our PAC estimates using the same approach as for coherence, and tested for significant PAC using a non-parametric cluster-based permutation test^80^.

### Stimulation during verbal free-recall memory task

Stimulation was applied only after a neurologist determined safe amplitudes using an iterative mapping procedure, stepping up stimulation in 0.5 mA increments and monitoring for after-discharges. The maximum amplitude selected (1.5 mA) fell well below standard safety boundaries for charge density^81^. We applied stimulation in a bipolar configuration, with current passing through a single pair of adjacent electrodes. Stimulation consisted of charge-balance biphasic rectangular pulses (width = 300 *μ*s) applied continuously at 50 Hz frequency for 4.6 s while subjects encoded two consecutive words. Then, stimulation was paused for the following two words, and then applied again, for each list of 12 words. Stimulation began 200 ms prior to word presentation and lasted until 200–450 ms after the offset of the second word in the stimulated pair. We applied stimulation during 20/25 of the lists in a session, so subjects were stimulated during encoding for 120 words and not stimulated for 180 words per session. To compare the role that stimulation played in predicting recall performance as a function of emotional context, we utilized weighted logistics regression models^82^, tested against each other using likelihood ratio tests.

Directly stimulating the brain while simultaneously recording iEEG signals often results in the appearance of artifactual signals while stimulation is delivered and following stimulation offset. In order to measure true physiological signals before and after stimulation, we followed prior methods in implementing an artifact detection algorithm to identify trials and channels to exclude from analysis with either complete signal saturation as well as gradual post-stimulation artifact^49,83^.

### Depression ratings

A subset of subjects performed the Beck Depression Inventory II, a self-assessment rating scale for depressive symptoms^84^. We utilized the conventional scoring criteria to categorize subjects as depressed or not depressed, as well as for the further categorization of depression severity. We excluded subjects with BDI-II scores between 5 and 13 to match the number of subjects with depression (n=22) to those without depression (n=24).

## Acknowledgements

We are grateful to the patients for participating in our study. This work was supported by NIH grant 2R01-MH104606 (to J.J.). We thank Molly Hermiller and Lukas Kunz for helpful comments and suggestions. We thank Mike Kahana for help with data collection.

## Author Contributions

S.E.Q. conceived the study; S.E.Q. and U.R.M. analyzed the data; J.M.S. processed neuroimaging data; and S.E.Q. and J.J. wrote the manuscript.

---

**Figure S1:**
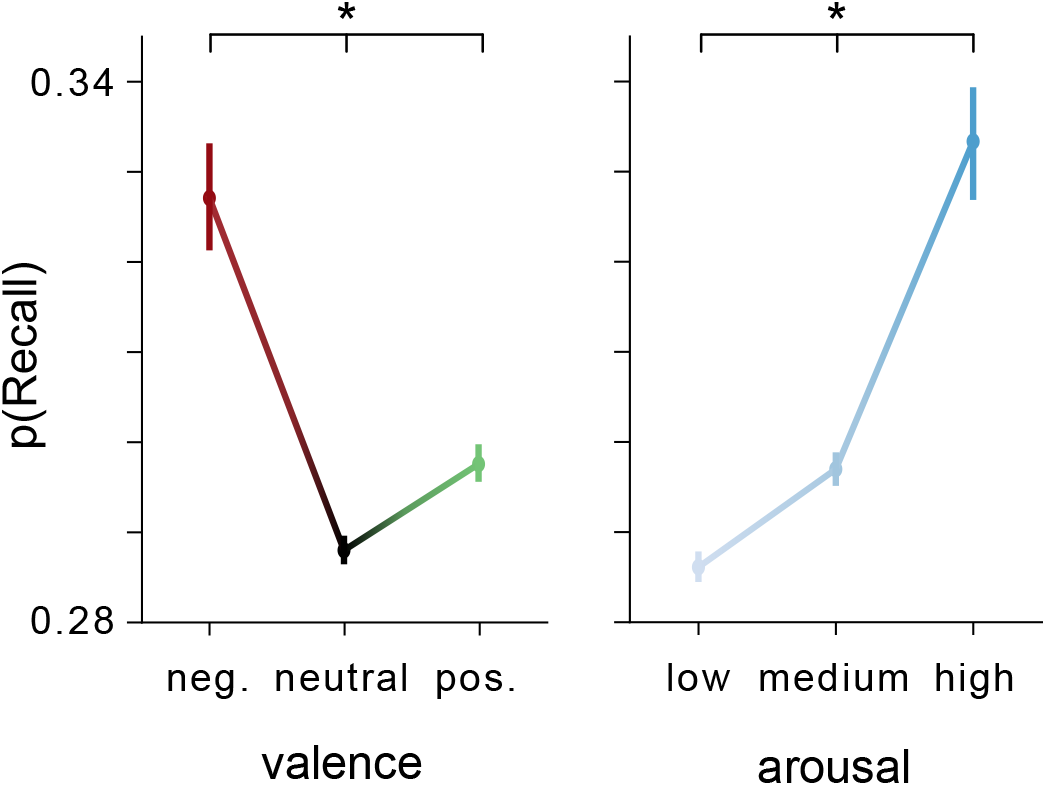
Effects of valence and arousal on recall are robust using a different ratings scale. Probability of recall significantly differed as a function of valence (*χ*(2) = 61, p< 8 × 10^-14^) and arousal (*χ*^2^(2) = 79, p< 7 × 10^-18^). Vertical bars denote standard error. Asterisks denote significant difference in proportions across categories. Related to Figure 1.

**Figure S2:**
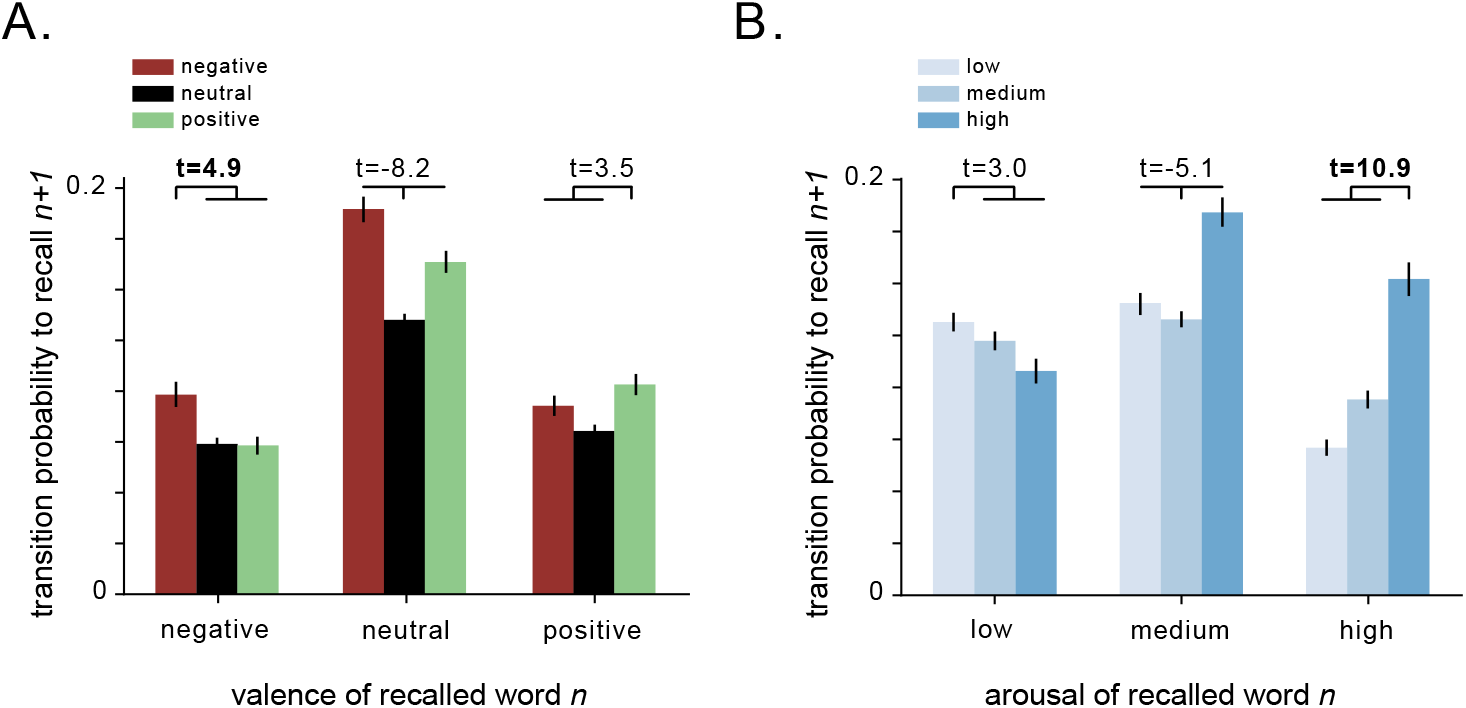
Emotional context modulates recall dynamics. A) Conditional response probability based on valence. The height of each bar depicts the probability of making a transition to a particular valence word (denoted by the color of the bar) as a function of the just recalled word’s valence (denoted by the x-axis label). Error bars denote standard error. T-statistics denote the relative proportion of within-valence transitions versus across-valence transitions. B) Conditional response probability based on arousal. The height of each bar depicts the probability of making a transition to a particular arousal word (denoted by the color of the bar) as a function of the just recalled word’s arousal (denoted by the x-axis label). Error bars denote standard error. T-statistics denote the relative proportion of within-arousal transitions versus across-arousal transitions. The largest t-statistic across both A and B is bolded. Related to Figure 1.

**Figure S3:**
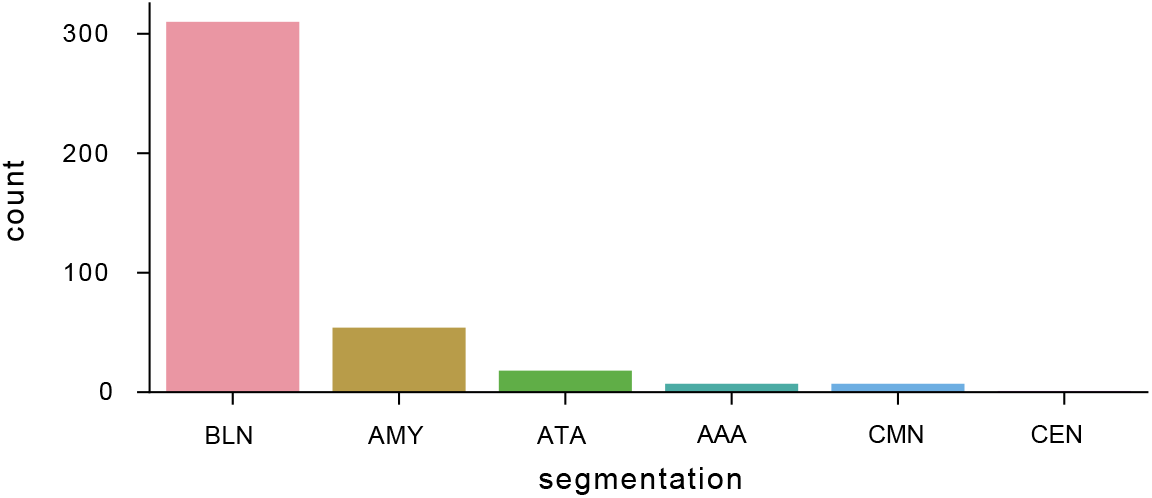
Segmentation of electrodes to different amygdala nuclei. Count of electrodes categorized to different amygdala nuclei on the basis of post-implant imaging. BLN = basolateral nuclei, ATA = amgydala transition areas, AAA = anterior amygdala area, CMN = cortical and medial nuclei, CEN = central nucleus, AMY = could not be localized to specific subregion. Related to Figure 2.

**Figure S4:**
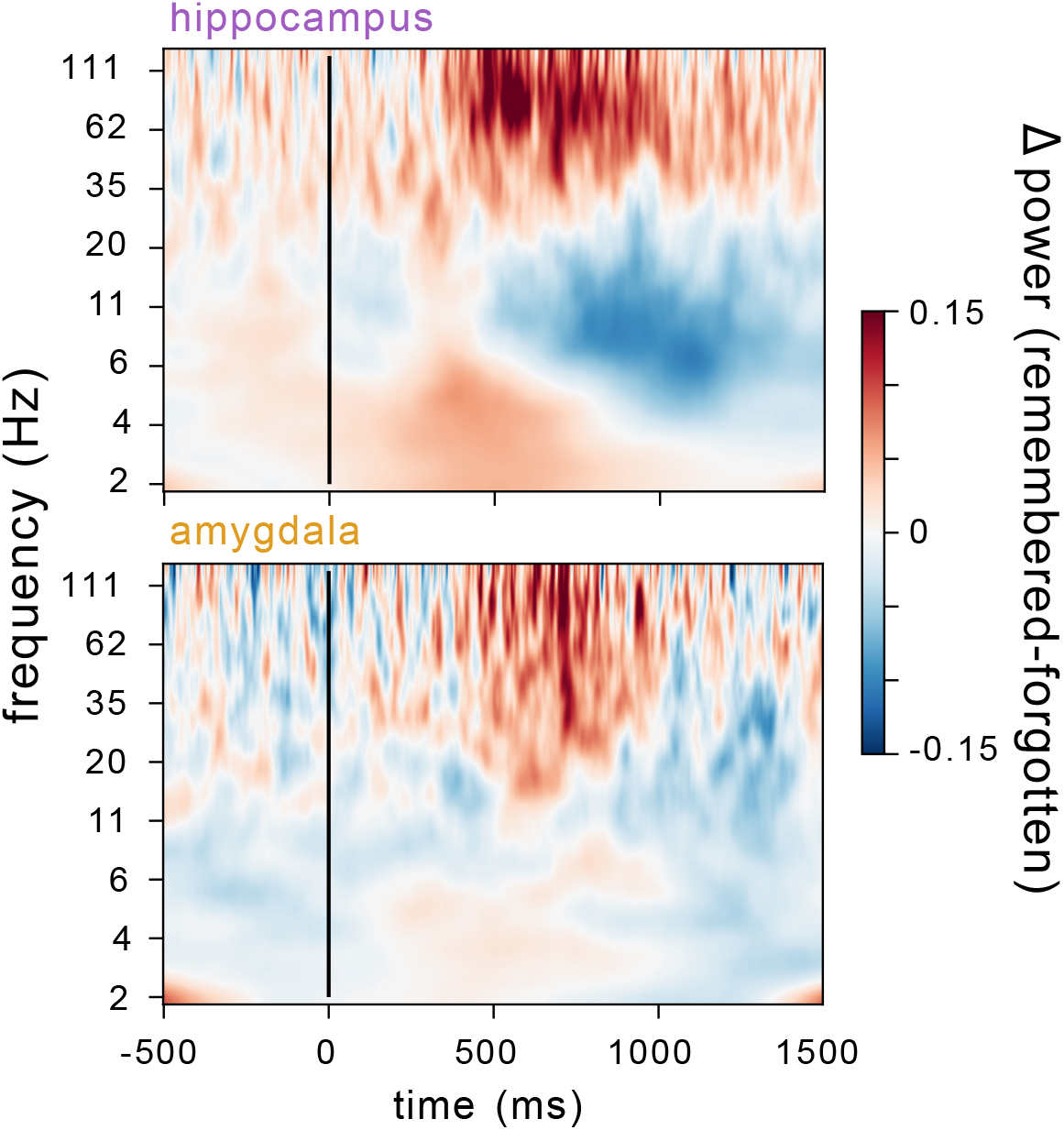
Hippocampal and amygdalar spectrogram depicting difference in power between remembered and forgotten trials across all electrodes. Average z-scored spectrogram for hippocampal (top) and amygdalar (bottom) electrodes showing difference between remembered and forgotten words. Warm colors indicate an increase in power during encoding of remembered words, while cool colors indicate a decrease in power. Vertical black line denotes the onset of word presentation. Related to Figure 2.

**Figure S5:**
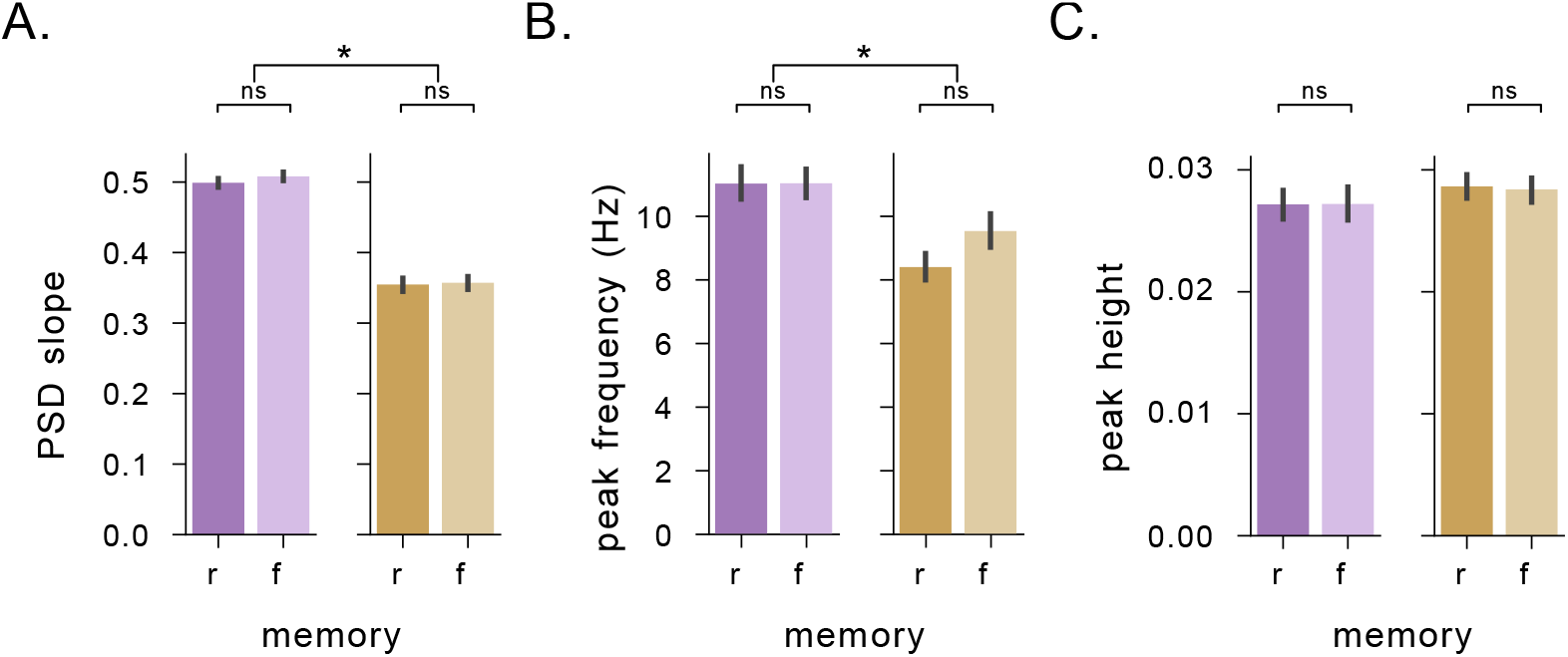
Memory-related power changes are not due to changes in spectra characteristics. A) Power spectra slope across the entire session for both remembered (dark shade) and forgotten (light shade) trials in both hippocampus (purple) and amygdala (orange). Asterisks denote significant differences (p’s< 7 × 10^-6^, t-test). B) Peak frequency across the entire session for both remembered and forgotten trials in both hippocampus and amygdala. C) Peak height across the entire session for both remembered and forgotten trials in both hippocampus and amygdala. Related to Figure 2.

**Figure S6:**
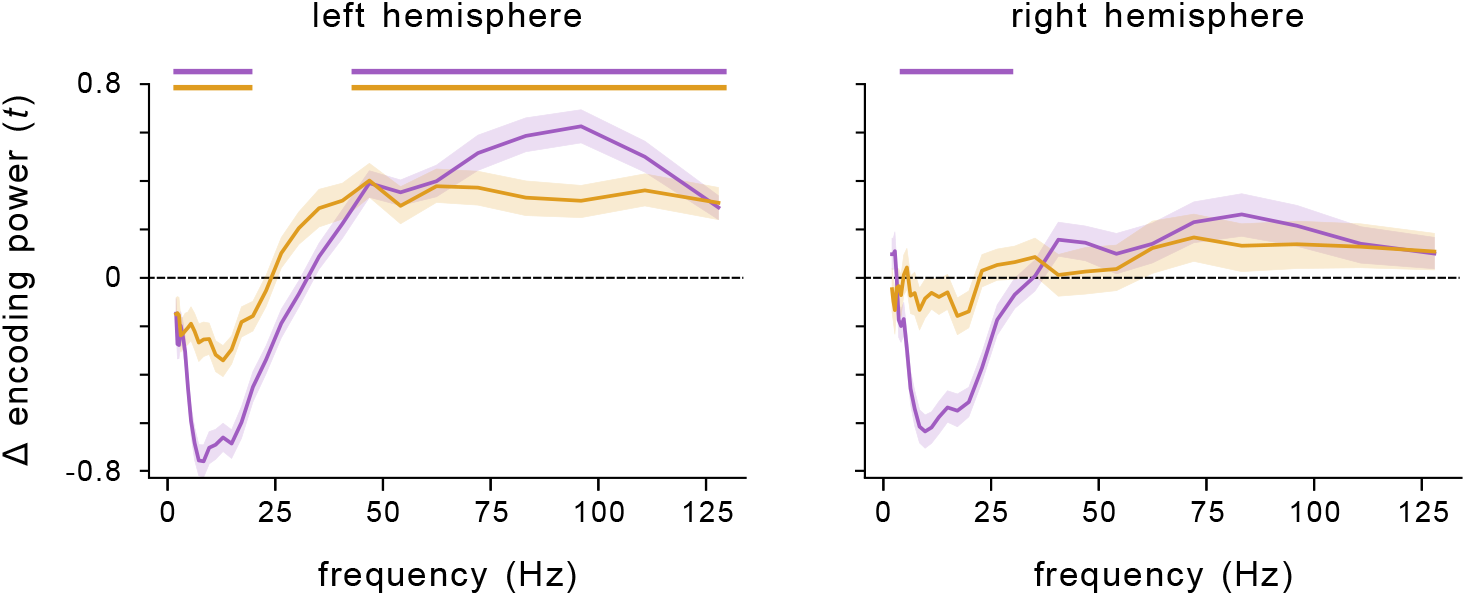
Laterality of HFA SME across all words. Comparison of SME (t-statistic) averaged over hippocampal (purple) and amygdala (orange) electrodes, separated by left and right hemispheres. Horizontal lines denote SMEs that significantly deviate from 0 (p’s< 0.05, cluster-permutation test). Related to Figure 2.

**Figure S7:**
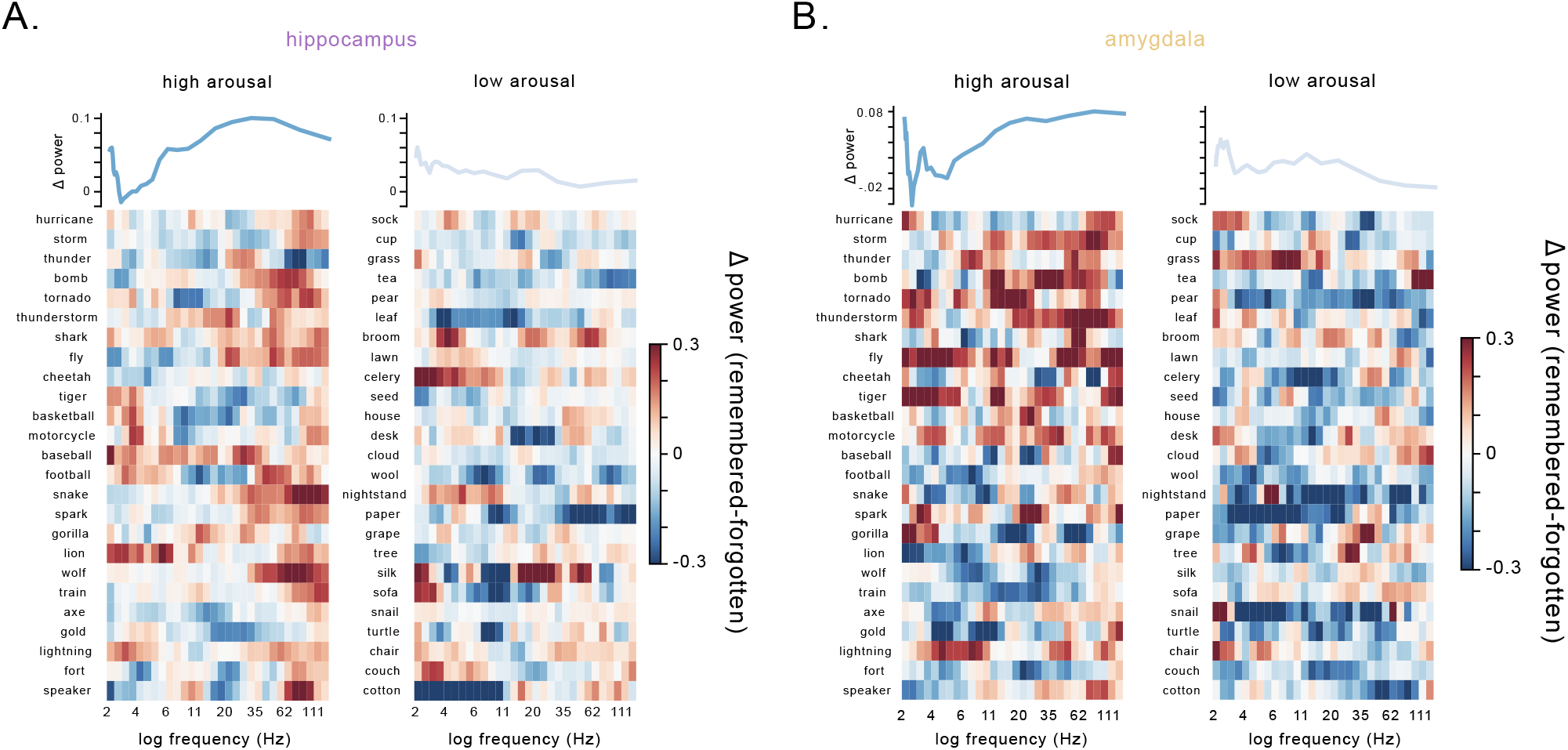
Word-level SME for high arousal and low arousal words averaged across the population. A) Heatmaps of hippocampal power (z-scored within session) for specific words from the task wordpool, averaged across sessions and subjects. Words were selected from the 30 words with the highest arousal ratings (left) or lowest arousal ratings (right). Warm colors indicate higher values while cool colors indicate lower values. Above each heatmap is the averaged z-scored power across the words in the heatmap. B) Same as panel A), but for amygdalar power. Related to Figure 2.

**Figure S8:**
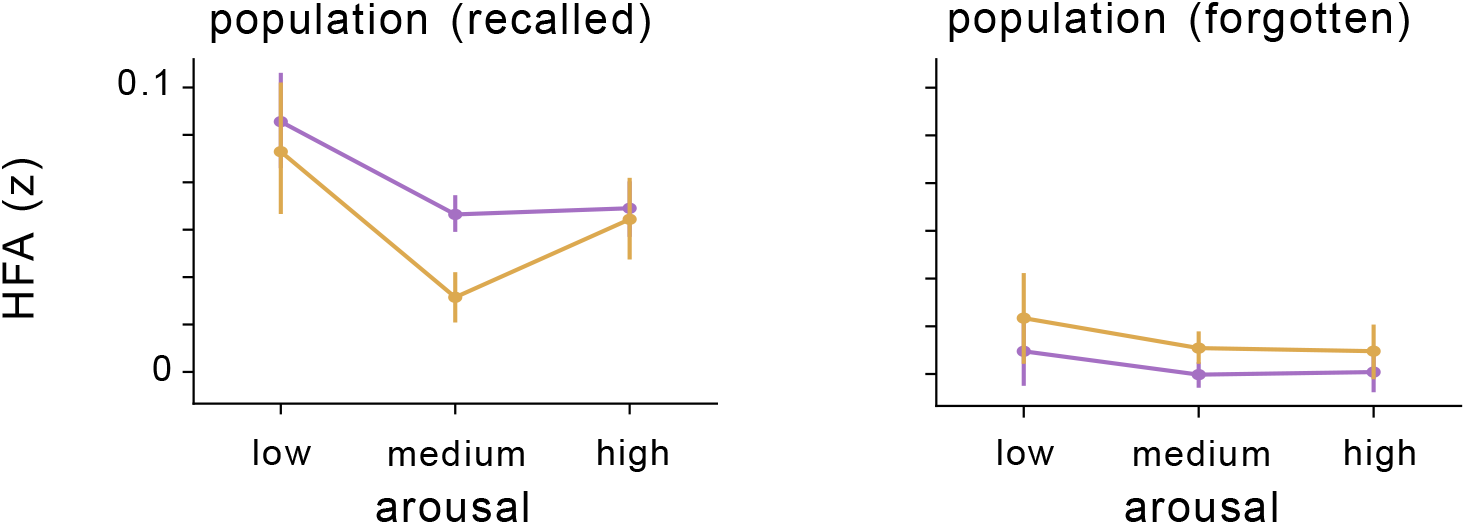
HFA SME as a function of valence across the population. Z-scored HFA in the amygdala (orange) and hippocampus (purple) during the encoding phase as a function of word valence for recalled (left) and forgotten (right) words. Circles represent mean of binned valence, with vertical lines denoting the standard error of the bin. Related to Figure 2.

**Figure S9:**
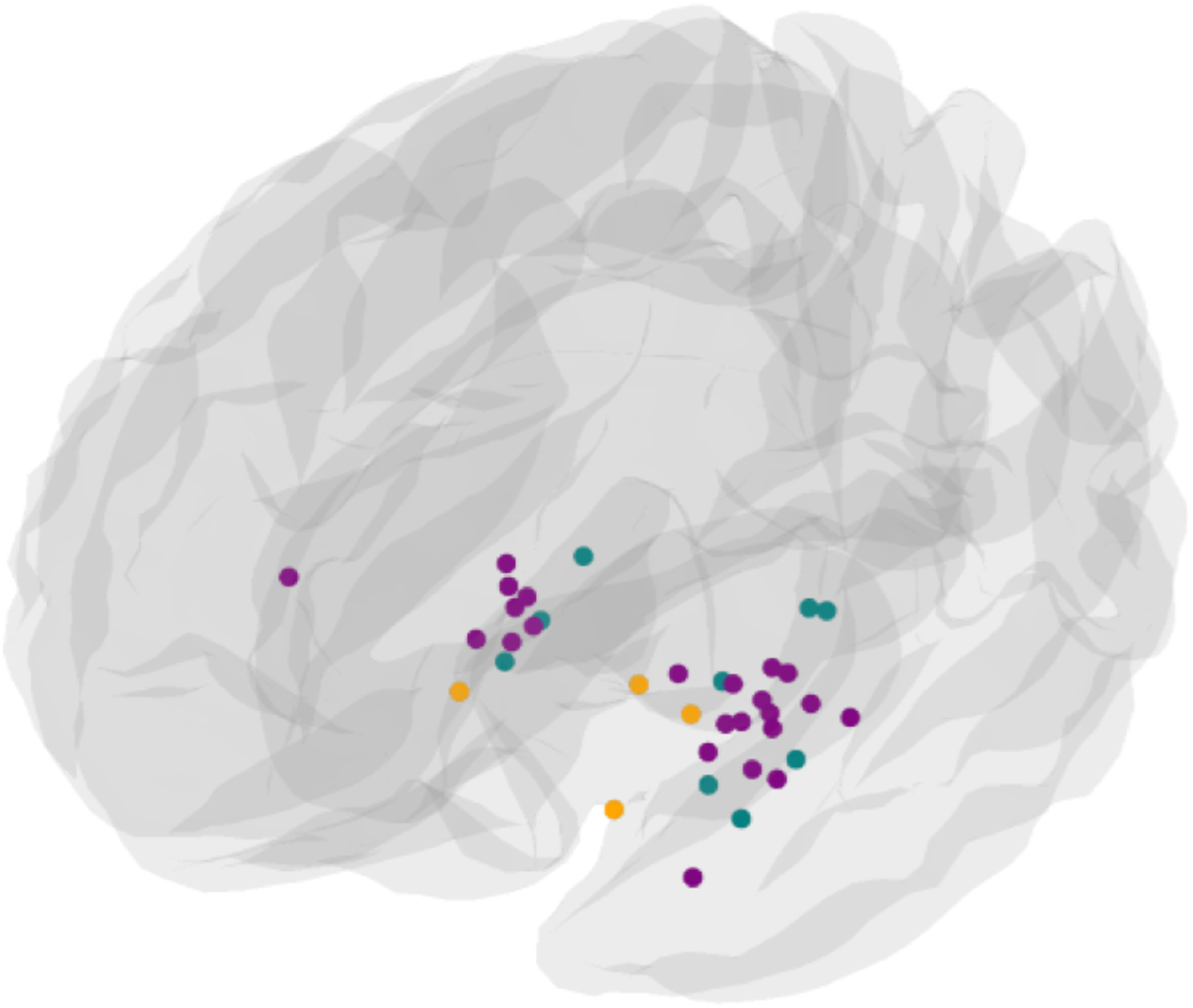
Location of stimulation electrodes. Hippocampal electrodes (purple), amygdala electrodes (orange), and non-hippocampal MTL electrodes (teal) where direct stimulation was applied. Related to Figure 3.

**Figure S10:**
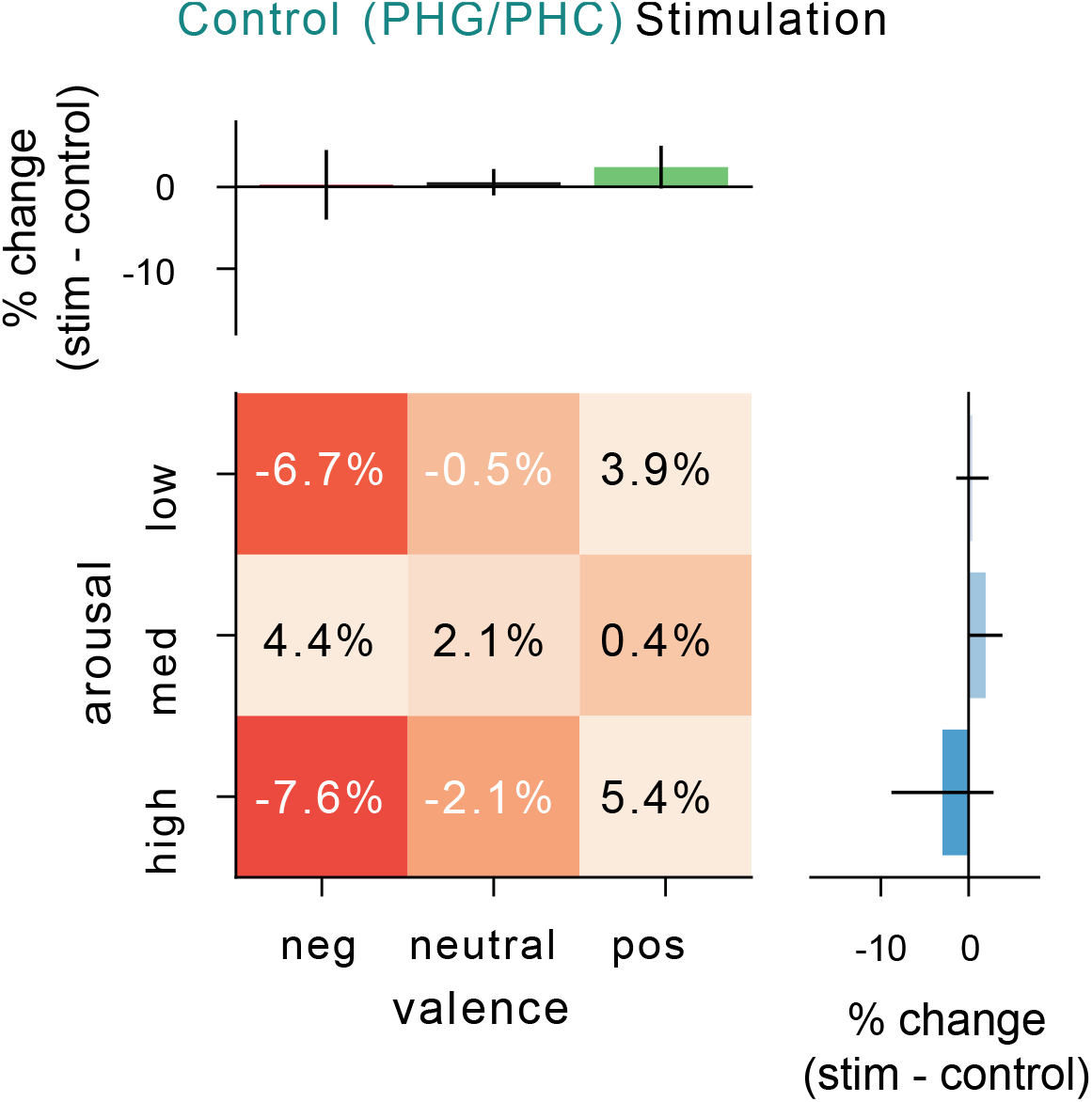
Stimulation in control regions in MTL does not result in significant modulation of memory as a function of emotional context. Effect of stimulation administered to electrodes located in the non-hippocampal MTL regions, split by arousal and valence. Asterisks indicate significant differences between conditions. Heatmap numbers indicate percentage of change in recall performance during stimulation. Related to Figure 3.

**Figure S11:**
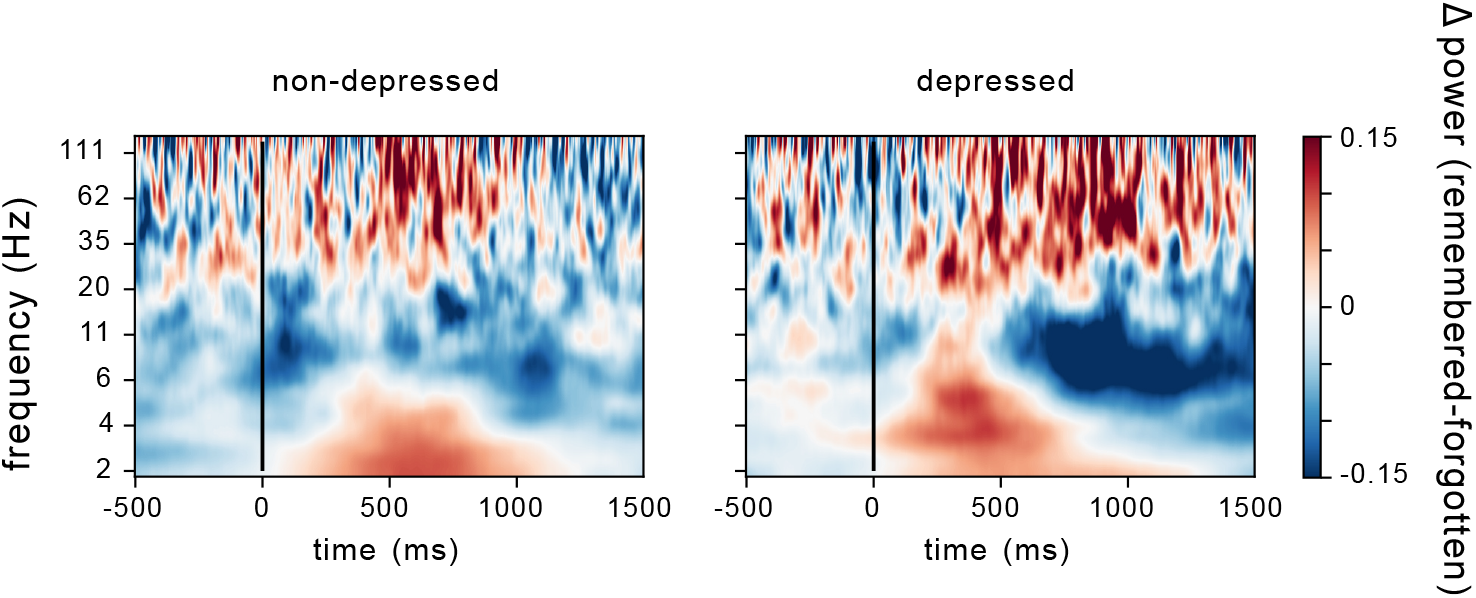
Difference in hippocampal SME between depressed and non-depressed patients. Average z-scored spectrogram for hippocampal electrodes in non-depressed (left) and depressed (right) subject showing difference between remembered and forgotten words. Warm colors indicate an increase in power during encoding of remembered words, while cool colors indicate a decrease in power. Vertical black line denotes the onset of word presentation. Related to Figure 4.

## References

[1] Bradley, M. M., Greenwald, M. K., Petry, M. C. & Lang, P. J. Remembering pictures: pleasure and arousal in memory. Journal of Experimental Psychology: Learning, Memory, and Cognition 18, 379 (1992).

[2] Kensinger, E. A. & Corkin, S. Memory enhancement for emotional words: are emotional words more vividly remembered than neutral words? Mem Cognit 31, 1169–80 (2003).

[3] Reisberg, D. & Hertel, P. Memory and emotion (Oxford University Press, Oxford, 2004). URL http://www.loc.gov/catdir/toc/ecip041/2003006595.html.

[4] Klein-Koerkamp, Y., Baciu, M. & Hot, P. Preserved and impaired emotional memory in alzheimer’s disease. Front Psychol 3, 331 (2012).

[5] Klüver, H. & Bucy, P. C. Preliminary analysis of functions of the temporal lobes in monkeys. J Neuropsychiatry Clin Neurosci 9, 606–20 (1939).

[6] Weiskrantz, L. Behavioral changes associated with ablation of the amygdaloid complex in monkeys. J Comp Physiol Psychol 49, 381–91 (1956).

[7] Poulin, S. P. et al. Amygdala atrophy is prominent in early alzheimer’s disease and relates to symptom severity. Psychiatry Res 194, 7–13 (2011).

[8] Tulving, E. & Markowitsch, H. J. Episodic and declarative memory: Role of the hippocampus. Hippocampus 8, 198–204 (1998).

[9] Adolphs, R., Cahill, L., Schul, R. & Babinsky, R. Impaired declarative memory for emotional material following bilateral amygdala damage in humans. Learn Mem 4, 291–300 (1997).

[10] LaBar, K. & Phelps, E. Arousal-mediated memory consolidation: Role of the medial temporal lobe in humans. Psychological Science 9, 490–493 (1998). URL https://doi.org/10.1111/1467-9280.00090. https://doi.org/10.1111/1467-9280.00090.

[11] Hamann, S. B., Ely, T. D., Grafton, S. T. & Kilts, C. D. Amygdala activity related to enhanced memory for pleasant and aversive stimuli. Nat Neurosci 2, 289–93 (1999).

[12] Canli, T., Zhao, Z., Brewer, J., Gabrieli, J. D. & Cahill, L. Event-related activation in the human amygdala associates with later memory for individual emotional experience. J Neurosci 20, RC99 (2000).

[13] Anderson, A. K. & Phelps, E. A. Lesions of the human amygdala impair enhanced perception of emotionally salient events. Nature 411, 305–9 (2001).

[14] Sharot, T. & Phelps, E. A. How arousal modulates memory: disentangling the effects of attention and retention. Cogn Affect Behav Neurosci 4, 294–306 (2004).

[15] Dolcos, F., LaBar, K. & Cabeza, R. Interaction between the amygdala and the medial temporal lobe memory system predicts better memory for emotional events. Neuron 42, 855–863 (2004).

[16] Phelps, E. A. Emotion and cognition: insights from studies of the human amygdala. Annu Rev Psychol 57, 27–53 (2006).

[17] Ahs, F., Kumlien, E. & Fredrikson, M. Arousal enhanced memory retention is eliminated following temporal lobe resection. Brain Cogn 73, 176–9 (2010).

[18] McGaugh, J. L. The amygdala modulates the consolidation of memories of emotionally arousing experiences. Annual Review of Neuroscience 27, 1–28 (2004).

[19] Atucha, E. et al. Noradrenergic activation of the basolateral amygdala maintains hippocampus-dependent accuracy of remote memory. Proc Natl Acad Sci U S A 114, 9176–9181 (2017).

[20] Cahill, L. & Alkire, M. T. Epinephrine enhancement of human memory consolidation: interaction with arousal at encoding. Neurobiol Learn Mem 79, 194–8 (2003).

[21] Hurlemann, R. et al. Noradrenergic modulation of emotion-induced forgetting and remembering. J Neurosci 25, 6343–9 (2005).

[22] Cahill, L., Prins, B., Weber, M. & McGaugh, J. L. Beta-adrenergic activation and memory for emotional events. Nature 371, 702–4 (1994).

[23] van Stegeren, A. H. The role of the noradrenergic system in emotional memory. Acta Psychol (Amst) 127, 532–41 (2008).

[24] Hagena, H., Hansen, N. & Manahan-Vaughan, D. -adrenergic control of hippocampal function: Subserving the choreography of synaptic information storage and memory. Cereb Cortex 26, 1349–64 (2016).

[25] McCall, J. G. et al. Locus coeruleus to basolateral amygdala noradrenergic projections promote anxiety-like behavior. Elife 6 (2017).

[26] Bacon, T. J., Pickering, A. E. & Mellor, J. R. Noradrenaline release from locus coeruleus terminals in the hippocampus enhances excitation-spike coupling in ca1 pyramidal neurons via-adrenoceptors. Cereb Cortex 30, 6135–6151 (2020).

[27] Haggerty, D. C., Glykos, V., Adams, N. E. & Lebeau, F. E. N. Bidirectional modulation of hippocampal gamma (20-80 hz) frequency activity in vitro via alpha()- and beta()-adrenergic receptors (ar). Neuroscience 253, 142–54 (2013).

[28] Cape, E. G. & Jones, B. E. Differential modulation of high-frequency gamma-electroencephalogram activity and sleep-wake state by noradrenaline and serotonin microinjections into the region of cholinergic basalis neurons. J Neurosci 18, 2653–66 (1998).

[29] Ridout, N., Astell, A., Reid, I., Glen, T. & O’Carroll, R. Memory bias for emotional facial expressions in major depression. Cogn Emot 17, 101–122 (2003).

[30] Pringle, A., McCabe, C., Cowen, P. J. & Harmer, C. J. Antidepressant treatment and emotional processing: can we dissociate the roles of serotonin and noradrenaline? J Psychopharmacol 27, 719–31 (2013).

[31] Scangos, K. W. et al. Closed-loop neuromodulation in an individual with treatment-resistant depression. Nat Med 27, 1696–1700 (2021).

[32] Mather, M., Clewett, D., Sakaki, M. & Harley, C. W. Norepinephrine ignites local hotspots of neuronal excitation: How arousal amplifies selectivity in perception and memory. Behav Brain Sci 39, e200 (2016).

[33] Madan, C. R. Exploring word memorability: How well do different word properties explain item free-recall probability? Psychon Bull Rev (2020).

[34] Aka, A., Phan, T. D. & Kahana, M. J. Predicting recall of words and lists. J Exp Psychol Learn Mem Cogn (2020).

[35] Kleinsmith, L. J. & Kaplan, S. Paired-associate learning as a function of arousal and interpolated interval. J Exp Psychol 65, 190–3 (1963).

[36] Kucewicz, M. T. et al. Pupil size reflects successful encoding and recall of memory in humans. Sci Rep 8, 4949 (2018).

[37] Posner, J., Russell, J. A. & Peterson, B. S. The circumplex model of affect: an integrative approach to affective neuroscience, cognitive development, and psychopathology. Dev Psychopathol 17, 715–34 (2005).

[38] LaBar, K. S. & Cabeza, R. Cognitive neuroscience of emotional memory. Nature Reviews Neuroscience 7, 54–64 (2006).

[39] Schildkraut, J. J. The catecholamine hypothesis of affective disorders: a review of supporting evidence. 1965. J Neuropsychiatry Clin Neurosci 7, 524–33; discussion 523–4 (1965).

[40] Mohammad, S. M. Obtaining reliable human ratings of valence, arousal, and dominance for 20,000 english words. In Proceedings of The Annual Conference of the Association for Computational Linguistics (ACL) (Melbourne, Australia, 2018).

[41] Kramer, J. H. et al. Hippocampal volume and retention in alzheimer’s disease. J Int Neuropsychol Soc 10, 639–43 (2004).

[42] Paller, K. A. & Wagner, A. D. Observing the transformation of experience into memory. Trends in Cognitive Sciences 6, 93–102 (2002).

[43] Prince, S., Daselaar, S. & Cabeza, R. Neural Correlates of Relational Memory: Successful Encoding and Retrieval of Semantic and Perceptual Associations. Journal of Neuroscience 25, 1203–1210 (2005).

[44] Sederberg, P. B. et al. Hippocampal and neocortical gamma oscillations predict memory formation in humans. Cerebral Cortex 17, 1190–1196 (2007).

[45] Lega, B., Jacobs, J. & Kahana, M. Human hippocampal theta oscillations and the formation of episodic memories. Hippocampus 22, 748–761 (2012).

[46] Burke, J. F. et al. Human intracranial high-frequency activity maps episodic memory formation in space and time. NeuroImage 85, 834–843 (2014).

[47] Long, N. M., Burke, J. F. & Kahana, M. J. Subsequent memory effect in intracranial and scalp EEG. NeuroImage 84, 488–494 (2014).

[48] Berridge, C. W. & Waterhouse, B. D. The locus coeruleus-noradrenergic system: modulation of behavioral state and state-dependent cognitive processes. Brain Res Brain Res Rev 42, 33–84 (2003).

[49] Mohan, U. R. et al. The effects of direct brain stimulation in humans depend on frequency, amplitude, and white-matter proximity. Brain Stimul 13, 1183–1195 (2020).

[50] Jacobs, J. et al. Direct electrical stimulation of the human entorhinal region and hippocampus impairs memory. Neuron 92, 1–8 (2016).

[51] Ilsley, J. E., Moffoot, A. P. & O’Carroll, R. E. An analysis of memory dysfunction in major depression. J Affect Disord 35, 1–9 (1995).

[52] Zheng, J. et al. Amygdala-hippocampal dynamics during salient information processing. Nat Commun 8, 14413 (2017).

[53] Murty, V. P., Ritchey, M., Adcock, R. A. & LaBar, K. S. fmri studies of successful emotional memory encoding: A quantitative meta-analysis. Neuropsychologia 48, 3459–69 (2010).

[54] Inman, C. S. et al. Direct electrical stimulation of the amygdala enhances declarative memory in humans. Proc Natl Acad Sci U S A 115, 98–103 (2018).

[55] Kensinger, E. A. & Corkin, S. Two routes to emotional memory: distinct neural processes for valence and arousal. Proc Natl Acad Sci U S A 101, 3310–5 (2004).

[56] Bijanki, K. R. et al. Case report: stimulation of the right amygdala induces transient changes in affective bias. Brain Stimul 7, 690–3 (2014).

[57] Harmer, C. J. et al. Effect of acute antidepressant administration on negative affective bias in depressed patients. Am J Psychiatry 166, 1178–84 (2009).

[58] El Mansari, M. et al. Relevance of norepinephrine-dopamine interactions in the treatment of major depressive disorder. CNS Neurosci Ther 16, e1–17 (2010).

[59] Gire, D. H. & Schoppa, N. E. Long-term enhancement of synchronized oscillations by adrenergic receptor activation in the olfactory bulb. J Neurophysiol 99, 2021–5 (2008).

[60] Marzo, A., Totah, N. K., Neves, R. M., Logothetis, N. K. & Eschenko, O. Unilateral electrical stimulation of rat locus coeruleus elicits bilateral response of norepinephrine neurons and sustained activation of medial prefrontal cortex. J Neurophysiol 111, 2570–88 (2014).

[61] Hajós, M., Hoffmann, W. E., Robinson, D. D., Yu, J. H. & Hajós-Korcsok, É. Norepinephrine but not serotonin reuptake inhibitors enhance theta and gamma activity of the septo-hippocampal system. Neuropsychopharmacology 28, 857–864 (2003). URLhttps://doi.org/10.1038/sj.npp.1300116.

[62] Headley, D. B. & Weinberger, N. M. Fear conditioning enhances oscillations and their entrainment of neurons representing the conditioned stimulus. J Neurosci 33, 5705–17 (2013).

[63] Ul Haq, R. et al. Adrenergic modulation of sharp wave-ripple activity in rat hippocampal slices. Hippocampus 22, 516–33 (2012).

[64] Sakon, J. J. & Kahana, M. J. Hippocampal ripples signal contextually-mediated episodic recall. bioRxiv (2021). URL https://www.biorxiv.org/content/early/2021/06/07/2021.06.07.447409. https://www.biorxiv.org/content/early/2021/06/07/2021.06.07.447409.full.pdf.

[65] Buchanan, T. W., Etzel, J. A., Adolphs, R. & Tranel, D. The influence of autonomic arousal and semantic relatedness on memory for emotional words. Int J Psychophysiol 61, 26–33 (2006).

[66] Bennett, C., Liang, K. C. & McGaugh, J. L. Depletion of adrenal catecholamines alters the amnestic effect of amygdala stimulation. Behav Brain Res 15, 83–91 (1985).

[67] O’Dell, T. J., Connor, S. A., Guglietta, R. & Nguyen, P. V. -adrenergic receptor signaling and modulation of long-term potentiation in the mammalian hippocampus. Learn Mem 22, 461–71 (2015).

[68] Manns, J. R. & Bass, D. I. The amygdala and prioritization of declarative memories. Curr Dir Psychol Sci 25, 261–265 (2016).

[69] Liu, Y., Cui, L., Schwarz, M. K., Dong, Y. & Schlüter, O. M. Adrenergic gate release for spike timing-dependent synaptic potentiation. Neuron 93, 394–408 (2017).

[70] de Voogd, L. D., Fernández, G. & Hermans, E. J. Disentangling the roles of arousal and amygdala activation in emotional declarative memory. Soc Cogn Affect Neurosci 11, 1471–80 (2016).

[71] Dahl, M. J., Mather, M. & Werkle-Bergner, M. Noradrenergic modulation of rhythmic neural activity shapes selective attention. Trends Cogn Sci (2021).

[72] Friendly, M., Franklin, P. E., Hoffman, D. & Rubin, D. C. The Toronto Word Pool: Norms for imagery, concreteness, orthographic variables, and grammatical usage for 1,080 words. Behavior Research Methods and Instrumentation 14, 375–399 (1982).

[73] Weidemann, C. T. et al. Neural activity reveals interactions between episodic and semantic memory systems during retrieval. Journal of Experimental Psychology: General (in press).

[74] Warriner, A. B., Kuperman, V. & Brysbaert, M. Norms of valence, arousal, and dominance for 13,915 English lemmas. Behavior research methods 45, 1191–1207 (2013).

[75] Pauli, W. M., Nili, A. N. & Tyszka, J. M. A high-resolution probabilistic in vivo atlas of human subcortical brain nuclei. Sci Data 5, 180063 (2018).

[76] Rhone, A. E. et al. A human amygdala site that inhibits respiration and elicits apnea in pediatric epilepsy. JCI insight 5 (2020).

[77] Gelinas, J. N., Khodagholy, D., Thesen, T., Devinsky, O. & Buzsáki, G. Interictal epileptiform discharges induce hippocampal–cortical coupling in temporal lobe epilepsy. Nature medicine 22, 641 (2016).

[78] Donoghue, T. et al. Parameterizing neural power spectra into periodic and aperiodic components. Nat Neurosci 23, 1655–1665 (2020).

[79] Tort, A. et al. Dynamic cross-frequency couplings of local field potential oscillations in rat striatum and hippocampus during performance of a T-maze task. Proceedings of the National Academy of Sciences, USA 105, 20517 (2008).

[80] Maris, E. & Oostenveld, R. Nonparametric statistical testing of EEG- and MEG-data. Journal of Neuroscience Methods 164, 177–190 (2007).

[81] Shannon, R. V. A model of safe levels for electrical stimulation. Biomedical Engineering, IEEE Transactions on 39, 424–426 (1992).

[82] Ezzyat, Y. & Rizzuto, D. S. Direct brain stimulation during episodic memory. Current Opinion in Biomedical Engineering 8, 78–83 (2018).

[83] Solomon, E. A. et al. Medial temporal lobe functional connectivity predicts stimulation-induced theta power. Nat Commun 9, 4437 (2018).

[84] Beck, A. T., Steer, R. A., Ball, R. & Ranieri, W. Comparison of beck depression inventories -ia and -ii in psychiatric outpatients. J Pers Assess 67, 588–97 (1996).

